# Decoding Root Biogeography: Building Reduced Complexity Functional Rhizosphere Microbial Consortia

**DOI:** 10.1101/2023.06.12.544662

**Authors:** Mingfei Chen, Shwetha Acharya, Mon Oo Yee, Kristine Grace Cabugao, Romy Chakraborty

**Affiliations:** Department of Ecology, Earth & Environmental Sciences Area, Lawrence Berkeley National Laboratory, Berkeley, California 94720, USA

**Keywords:** rhizosphere, high-throughput, microbial, root exudates, reduced complexity consortia

## Abstract

The rhizosphere microbiome plays a crucial role in supporting plant productivity and contributes to ecosystem functioning by regulating nutrient cycling, soil integrity, and carbon storage. However, characterizing their functional attributes and microbial relationships remains challenging due to their complex taxonomic and functional compositions. To enable such studies, the development of reduced complexity microbial consortia derived from the rhizosphere microbiome of the natural ecosystem is highly desirable. Designing and assembling reduced complexity consortia that mimic natural communities with consistent, stable, predictable features are highly sought after but is challenging to deliver. Here we present our systematic controlled design towards successful assembly of several such rhizosphere derived reduced complexity consortia. From *Brachypodium* grown in natural soil under controlled lab conditions, we enriched the root-associated microbes, utilizing carbon compounds prevalent in Brachypodium root exudates. By transferring the enrichments every 3 or 7 days for 9 generations, we developed both fast and slow-growing microbial communities. 16S rRNA amplicon analysis revealed that both inoculum and carbon substrates significantly influence microbial community composition. For example, 1/10 R2A preferentially enriched Amplicon Sequence Variants (ASVs) from slow growing taxa vital to plant including Acidobacteria and Verrucomicrobia. Network analysis revealed that although fast and slow growing microbial consortia have distinct key taxa, the key hubs (keystone taxa) for both belong to genera with plant growth promoting (PGP) traits. This suggests that PGP bacteria might play a central role in controlling the microbial networks among rhizospheric microbiomes. Based on the stability and richness results from different transfers, most carbon substrates lead to microbial consortia with reduced complexity and high stability after a few transfers. The stability tests of the derived microbial consortia also showed high stability, reproducibility, and revivability of the constructed microbial consortia. Our study represents a significant step towards understanding and harnessing the potential of rhizosphere microbiomes, with implications for sustainable agriculture and environmental management.

## 1. Introduction

Rhizosphere microbes, localized immediately surrounding plant roots, have co-evolved with host plants to form mutually beneficial relationships. Rhizosphere microbes possess a range of functions, including plant growth promotion traits that benefit their host plants. These traits encompass the conversion of essential nutrients (e.g., nitrogen, phosphate, zinc, iron) into more accessible forms for plant assimilation [1–4], conferring resilience against environmental stressors such as pathogen infections and water limitations [5–8], and the secretion of plant growth promoting hormones [8, 9]. In return, rhizosphere microbes receive carbon from the root exudates of the plant [10–12]. Given their high abundance, diversity, and activity, rhizosphere microbes are considered as “the second genome of plants” [13] and act as key drivers of soil nutrient cycling and carbon sequestration [14, 15]. For the purpose of advancing our understanding and a more comprehensive study of their pivotal functions, it is critical to recover and to recreate a representative rhizosphere community in the laboratory setting that accurately encapsulates the inherent diversity and functions of the rhizobiome.

Prior studies indicate that within the rhizosphere, several microbial species work collectively within a community, promoting advantageous growth traits in the host plant. Two methods, known as bottom-up and top-down, are typically employed to construct synthetic communities that reflect native microbes. The bottom-up method focuses on building synthetic communities from isolates directly cultivated from the source. However, this method could exclude rare and/or dormant taxa [16–18] and demands pre-existing knowledge about microbial interactions for successful consortia assembly and stability [19, 21, 22]. Conversely, the top-down approach employs targeted enrichments such as limiting carbon sources or antibiotic treatments to generate consortia with reduced complexity [19, 20]. However, this approach could limit the phylogenetic diversity of the consortia by only recovering fast-growing generalists due to the use of easily degradable carbon sources or restricted variables [21, 22], and still requires a detailed analysis of individual microbial consortia members to understand their interactions within the community [21]. Consequently, there is a lack of effective strategies for creating robust, field-derived rhizosphere microbial consortia that demonstrate stability, complexity, reproducibility, scalability, and genetic tractability [21, 26].

To address this, we developed a novel, systematic, and standardized pipeline that integrates approaches to generate reduced complexity microbial consortia that optimally represent the rhizosphere microbiome of *Brachypodium distachyon* and satisfies all the above criteria. We first grew young *Brachypodium distachyon* using naturally occuring soil in standardized fabricated ecosystems called EcoFABs and controlled pots and tubes [24]. We demonstrated that the rhizosphere microbiome of *Brachypodium* was clearly distinct from microbiome of surrounding bulk soil [25], and that the microbial community in EcoFABs is highly reproducible [25]. In this study, we performed a large number of high-throughput enrichments to assess the effects of various critical parameters, such as the original rhizosphere inoculum, carbon substrate, and microbial growth rates, on the composition of reduced complexity microbial consortia. Subsequently, we identified microbial diversity and richness, the key microbial taxa significantly impacted by these factors and the keystone taxa that govern the interactions between the core community members. Furthermore, we demonstrated the validity of our pipeline showing stability, tractability, reproducibility, and revivability. Our research has the potential to significantly advance rhizosphere microbiome investigations by providing a robust and standardized pipeline for developing representative microbial consortia. The findings from this study may have far-reaching implications in agriculture, ecosystem management, and climate change mitigation strategies.

## 2. Materials and Methods

### Microbial community inocula

The enrichment inoculum consisted of rhizosphere soil from *Brachypodium distachyon* grown in soil collected from our field site in Angelo Coast Range Reserve, California (39° 44′ 21.4′′ N 123° 37′ 51.0′′ W) and in three types of containers-fabricated ecosystems (EcoFABs), pots and test tubes [25]. The glycerol stocks containing the loosely bound and tightly bound soil from the base and tip of the 14-day old *B. distachyon* roots were thawed and combined for each container type in order to prepare enrichment inocula. Inoculum for each container type was normalized to 4.6 x 10^6^ cells ml^-1^ in RCH2 media and 10% of these resulting soil slurries (180 μl) was used to inoculate enrichment media.

### Media for enrichment

The base media for most enrichments was a modified RCH2 media at pH 6.0 [26](Supplementary Table S1). After autoclaving, we added 10 ml/L of Wolfe’s vitamins [27]. Carbon source stocks were prepared in milliQ water at 20 mM, filter (0.2 μm) sterilized, and added to sterile RCH2 media at a final concentration of 2 mM. For carbon sources, single carbon sources relevant to *Brachypodium distachyon* root exudates were used: citrate, malate, succinate, salicylic acid, fumaric acid, glucose, sucrose, arabinose, asparagine, glycine, glutamine, serine, galacturonic acid, glucuronic acid, and a mixture of all these carbon sources (also at 2 mM). 2 mM salicylic acid was prepared in RCH2 media since it was not soluble in milliQ water at higher concentrations. Lastly, 1/10th dilution of commercially available media, R2A (hereafter referred to as 1/10 R2A) was used as a reference since R2A has been widely adopted for enrichment and isolation of soil and subsurface microbes [21, 28–31].

### High-throughput enrichment and sampling

Targeted, high-throughput (HT) enrichments via sequential transfers in 96-well plates were performed to get reduced complexity communities. Each well had 1.8 ml volume in total with 10% inoculum (180 μl) which originates from *Brachypodium distachyon* rhizosphere samples grown in different containers (EcoFABs, tubes, pots) and medium with either 2mM carbon source in RCH2 or 1/10 R2A. In each transfer, there were two plates where each row contains a different carbon source, and columns containing inoculum replicates from different containers (Supplementary Table S2). Each set (A and B) were transferred 10 times at two different intervals, 3 days and 7 days. Each transfer is referred to as a “Generation”, which indicates how many transfers have occurred (e.g., Gen 6 means it is the 6th transfer from the original plate). Plates were incubated in the 30 °C shaking incubator at 130 rpm, after sealing with a breath-easy seal (Parafilm). For every 3 or 7 days, 180 μl (10%) of inoculum was transferred to new plates with fresh media. Glycerol stocks (80 μl inoculum + 40 μl glycerol) were prepared each transfer and frozen at −80 °C. Ultimately, 40 plates containing 3840 samples were obtained during the enrichment process.

### Stability test

In order to assess microbial stability/volatility over time, the volatility analysis was implemented as previously described [32], in which the amount of the community change between successive time points was measured with weighted UniFrac distances calculated from the ‘rbiom’ package. The reproducibility and revivability of our derived microbial consortia were further assessed using five glycerol stocks (3-day: 3B8-H4, 3B8-H11; 7-day: 7B5-H1, 7B5-H8, 7B5-H11) from our enrichment experiments with distinct microbial communities. The naming convention (e.g., “3B8-H4”) indicates a 3-day or 7-day time interval, generation 8 or 5, and the specific location of a glycerol stock in a 96 deep well plate. These glycerol stocks were selected considering the microbial diversity observed in the enrichment experiments, while also incorporating various time intervals. Glycerol stocks were completely thawed (120 uL) and serially subcultured (10% inoculum and 90% media) to bring the final volume up to 100 mL until OD600 reached 0.4. The culture was then spun down as pellets for DNA extraction (80 mL), saved as 10 glycerol stocks (1 mL culture + 0.5 mL glycerol), and used for the next round of subculture (10 mL). The reproducibility of our consortia was tested by continuing subculturing using a 10% v/v inoculum every 3 or 7 days, totaling 5 transfers. For revivability tests, the preserved glycerol stocks from the first round of subculture were thawed, the volumes were brought up to 100 mL, and the pellets were spun down for DNA extraction (80 mL).

### 16S rRNA gene amplicon data processing and amplicon sequence variants analysis

Genomic DNA of four timepoints (transfers 1, 3, 6 and 9) and eight carbon sources (glucose, citrate, glutamine, asparagine, glucuronic acid, mixed carbon, and 1/10 R2A) from enrichment experiments and six transfers of tested samples from stability tests were extracted at UC Berkeley DNA Sequencing Facility and submitted to Novogene Corporation Inc. for 16S amplicon sequencing of the V4 region using the universal bacterial primers 515F (GTGCCAGCMGCCGCGGTAA) and 806R (GGACTACHVGGGTWTCTAAT). The method previously published [21] was employed to analyze the amplicon sequence variants (ASVs). In summary, analysis of microbial community sequences was conducted using QIIME 2 2020.8 [33]. Novogene provided demultiplexed sequences that were imported using the Phred33V2 variant. Quality filtering and denoising steps were performed with DADA2 [34] and amplicon sequence variants were aligned with MAFFT [35]. Taxonomy was assigned using the Silva reference database v138 from 515F/806R region of sequences classifier, using a cutoff at 99% for ASVs [36–39]. The phylogenetic tree of enriched ASVs was constructed with RAxML [40], and the tree visualization was performed using iTOL [41].

### Data analysis and statistics

Shannon’s diversity index (H’) and multivariate statistics were performed using the R package ‘vegan’ [42]. ASV distributions were transformed into relative abundances using the function ‘decostand’. These were subjected to Hellinger transformation before calculation of Bray-Curtis dissimilarity matrices comparing community composition between samples. Principal coordinate analysis (PCoA) using the function ‘vegdist’ was performed using these dissimilarity matrices. A permutational multivariate analysis of variance (PERMANOVA) model was implemented in the vegan function ‘adonis’ using Bray-Curtis distance matrix to evaluate the effect of C source, original inoculum, and transfer interval on community structure. The relative abundance of taxa among the samples were compared and enriched ASVs were selected using DESeq2 packages from the R software [43].

To understand the interactions between different species under different treatments, core ASVs (present in >75% samples from the 4 generations) from the enrichment experiments were selected and their abundance matrix from all samples was loaded into R to construct and visualize the network using the “NetCoMi” package [44]. Pearson correlation with a threshold value > 0.3 and student t-test with a corresponding p-value of < 0.05 were used to generate the sparse matrix for the network analysis. In this co-occurrence network, each node represents a single ASV. The edges connecting the nodes indicate a strong and significant correlation between the ASVs. The clustering in this network was used to group the ASVs into modules that were densely connected within themselves but sparsely connected to other modules.

## 3. Results

### Carbon sources and original inocula shape the diversity of enrichments

All variables tested in the enrichment experiments (C sources, original inoculum, transfer interval) significantly influenced bacterial community structure in our enrichments. PERMANOVA results considering all parameters show that original inoculum (*R*^2^ = 0.160, *p* < 0.001) and carbon sources (*R*^2^ = 0.149, *p* < 0.001) are the major drivers of community dissimilarity. Original inoculum means what? Describe here. Different sampling intervals (*R*^2^ = 0.030, *p* = 0.001) contributed to variation to a lesser extent. We then visualized the grouped samples by PCoA ordination comparing different variables (Figure 1A-C), and used pairwise PERMANOVA comparisons to validate their statistical significance (Supplementary Table S3). When comparing across different inocula, the microbial communities derived from EcoFAB inocula clearly differ from pot or tube inocula (Figure 1A). Upon evaluating the impact of carbon substrates, significant variations in microbial community composition were observed for all tested carbons, with those amended with 1/10 R2A markedly differing from all other carbon sources (Figure 1B). Enrichments transferred at 3-day or 7-day intervals showed no distinct differences solely based on observation of the PCoA plot, but still show significant impacts on communities from PERMANOVA results (Figure 1C).

**Figure 1:**
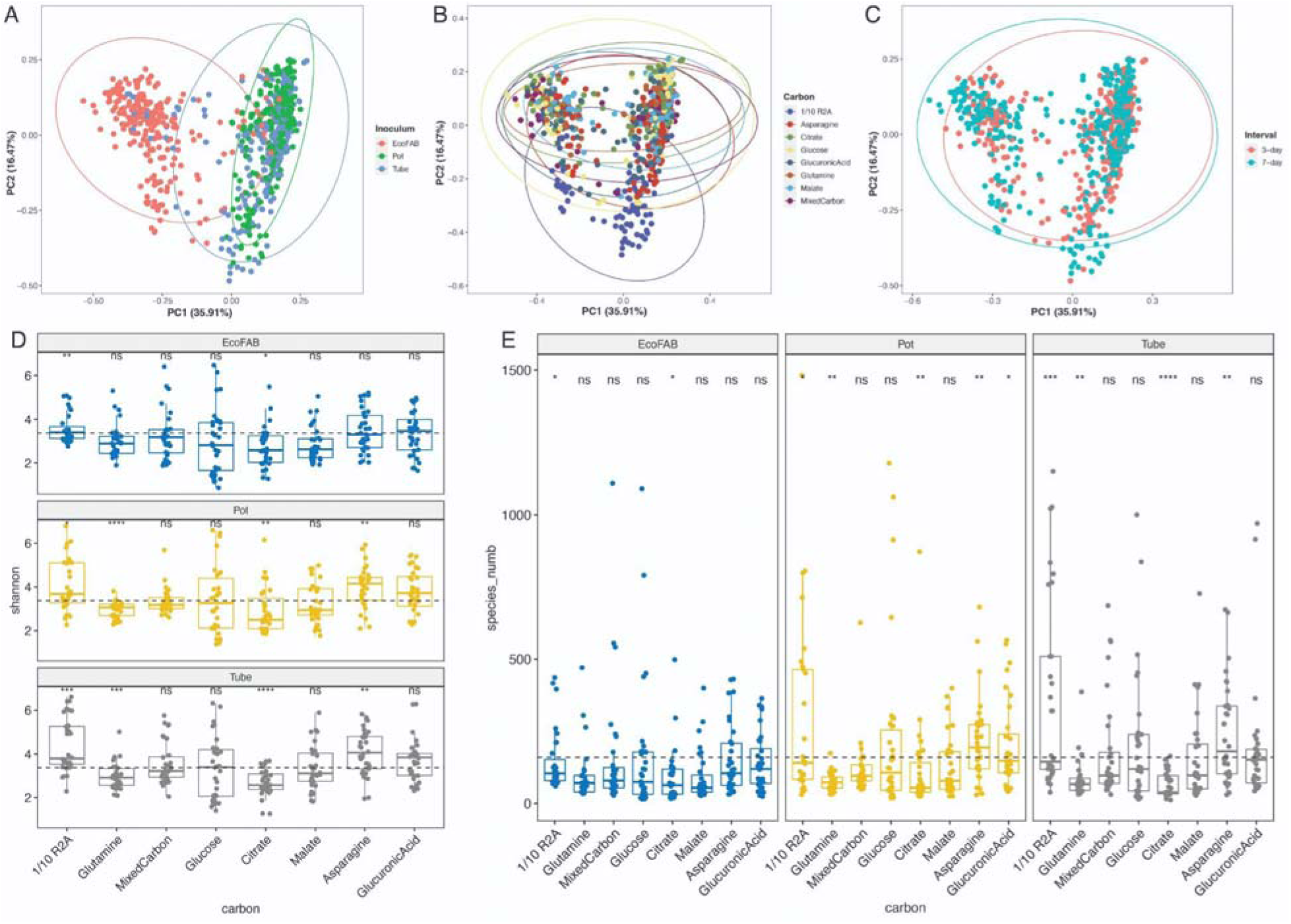
PCoA plot based on Bray-Curtis distance show enrichment results comparing (A) original inoculum, (B) amended carbon substrates, and (C) transfer intervals. Boxplot showing Shannon (D) and richness (E) diversity indexes of enrichment results, separating by different containers and amended carbon substrates. Significant differences compared to global mean values of shannon and richness indexes are indicated in asteroid (*). * =, ** =, *** =, **** =.

We also observed that carbon sources and the original inocula drive the biodiversity of enrichments. When compared against mean Shannon diversity index across all inocula, samples amended with citrate show significantly lower diversity values while samples amended with 1/10 R2A show significantly higher diversity values (Figure 1D-E). Asparagine can significantly increase the diversity indexes for pot and tube samples, while glutamine can decrease them. However, both amino acids show minimal impacts on the samples from the EcoFAB inocula (Figure 1D-E). Overall, enrichments derived from EcoFAB inoculum have significantly lower Shannon indices than pot and tube inoculum (Supplementary Figure S1A). With regards to different time intervals, 3-day samples have significantly higher diversity than the 7-day samples (Supplemental Figure S1B).

### Different carbon substrates can enrich unique rhizosphere taxa

Since we observed significant impact of carbon substrates on the microbial community composition, we then used differential gene relative abundance analysis (DESeq2) to quantitatively examine the impacts of carbon substrates on different taxa. We first examined which ASVs were significantly enriched in one carbon source over another. Given that original inoculum also showed significant impacts on the microbial communities (Figure 1), we separated the different original inoculum and compared the ASVs of different carbon sources respectively (Figure 2). We only include the abundant taxa with a baseMean value > 5 with a total of 93 enriched ASVs, after removing potential contaminants (i.e., abundant ASVs that only present in one sample). Among the 93 enriched ASVs, 48 of them show matches with zOTU data from prior biogeography experiments) [25] (97% cutoff (Supplementary Table S5). Generally, 1/10 R2A had the highest potential to enrich ASVs compared to other carbon sources (48 enriched ASVs, 28 matching ASVs). Some key taxa that were significantly enriched in 1/10 R2A across all three original inocula include ASVs from genera *Terriglobus*, *Castellaniella*, *Arthrobacter*, *Luteibacter*, *Bacillus* and unclassified Rhizobiaceae (Supplementary Table S5). Few other carbon sources also enriched unique genera/ASVs matching with zOTUs. For example, glucose significantly enriched ASVs from genera *Delftia* and *Mycobacterium*, glucuronic acid significantly enriched ASVs from genus *Romboutsia*, and glutamine significantly enriched ASVs from genus *Brevibacillus* (Supplementary Table S5). In addition, different carbon sources could also enrich different ASVs within the same genera. For example, asparagine, glucose, and mixed carbon all enriched distinct ASVs from the genus *Allorhizobium-Neorhizobium-Pararhizobium-Rhizobium*.

**Figure 2:**
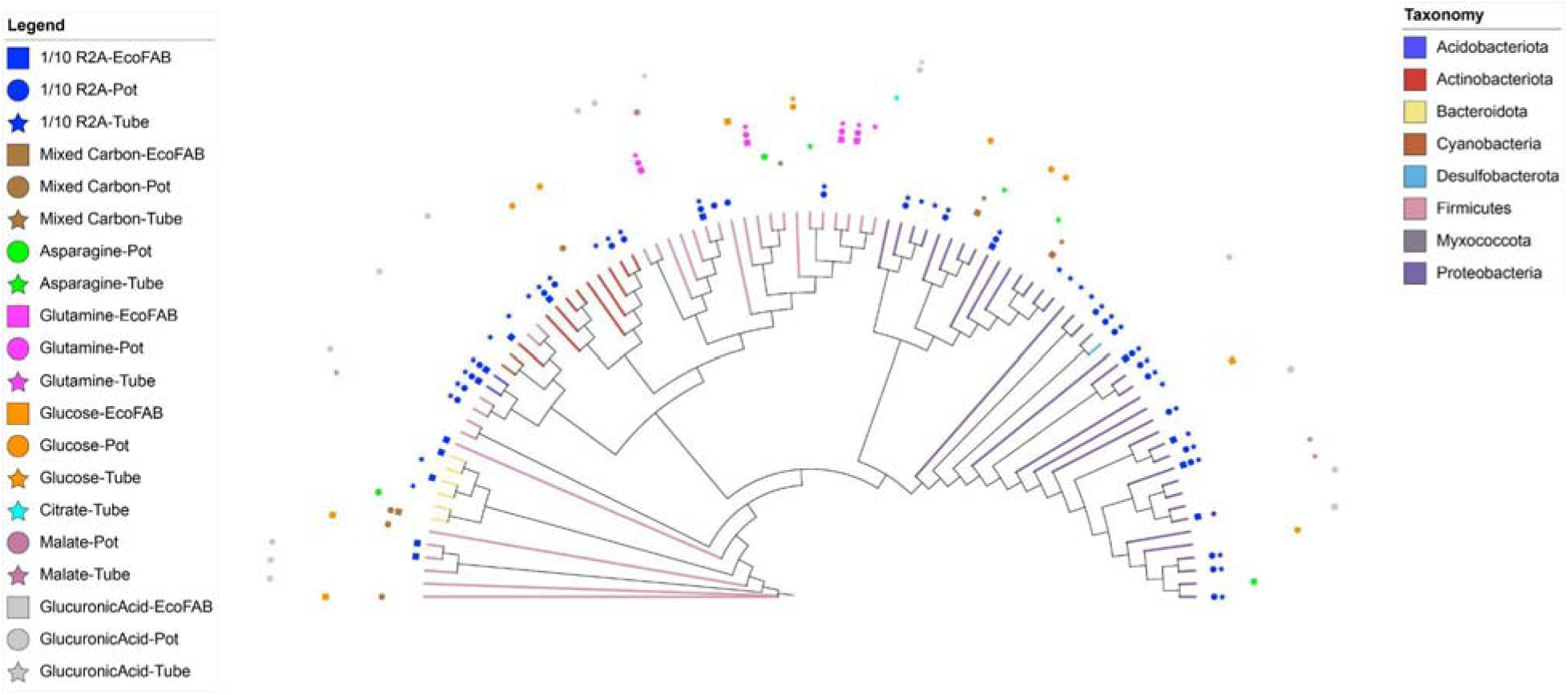
Neighbor-joining tree of 93 ASVs that are significantly enriched in one carbon substrate than others (DESeq2 results with p < 0.05 and rowMean values > 5). Markers surrounding the tree denote the original inoculum (EcoFAB - square, Pot - circle, or Tube - star) and C substrate in which the ASV was significantly enriched.

### Complex carbon sources can enrich slow-growing but important rhizosphere microbes

Next, we compared how different carbons can impact microbes with different growth rates. We classified microbes that were more abundant in 7-day samples as slow-growing microbes and those that were more abundant in 3-day samples as fast-growing microbes. We only included Gen 6 and Gen 9 samples for DESeq2 analysis because Gen 6 and Gen 9 were later stage transfers which contained very low amounts of soil-associated nutrients from the inoculum and would mainly utilize amended carbon substrates as carbon source. When comparing the results of slow-growing microbes and fast-growing microbes with different carbon sources (Figure 3A), we found that 1/10 R2A, followed by asparagine and glucuronic acid, had the highest potential to enrich both slow and fast growers, while citrate and glutamine had the least potential (Figure 3B-C). These results are consistent with Shannon diversity results comparing different carbon substrates (Figure 1D-E). In addition, different carbon sources exhibited differential enrichment of fast- and slow-growing microbes. 1/10 R2A, glucuronic acid and asparagine substantially enriched slow-growing microbes (Figure 3B), while glucose and malate substantially enriched fast-growing microbes (Figure 3C).

**Figure 3:**
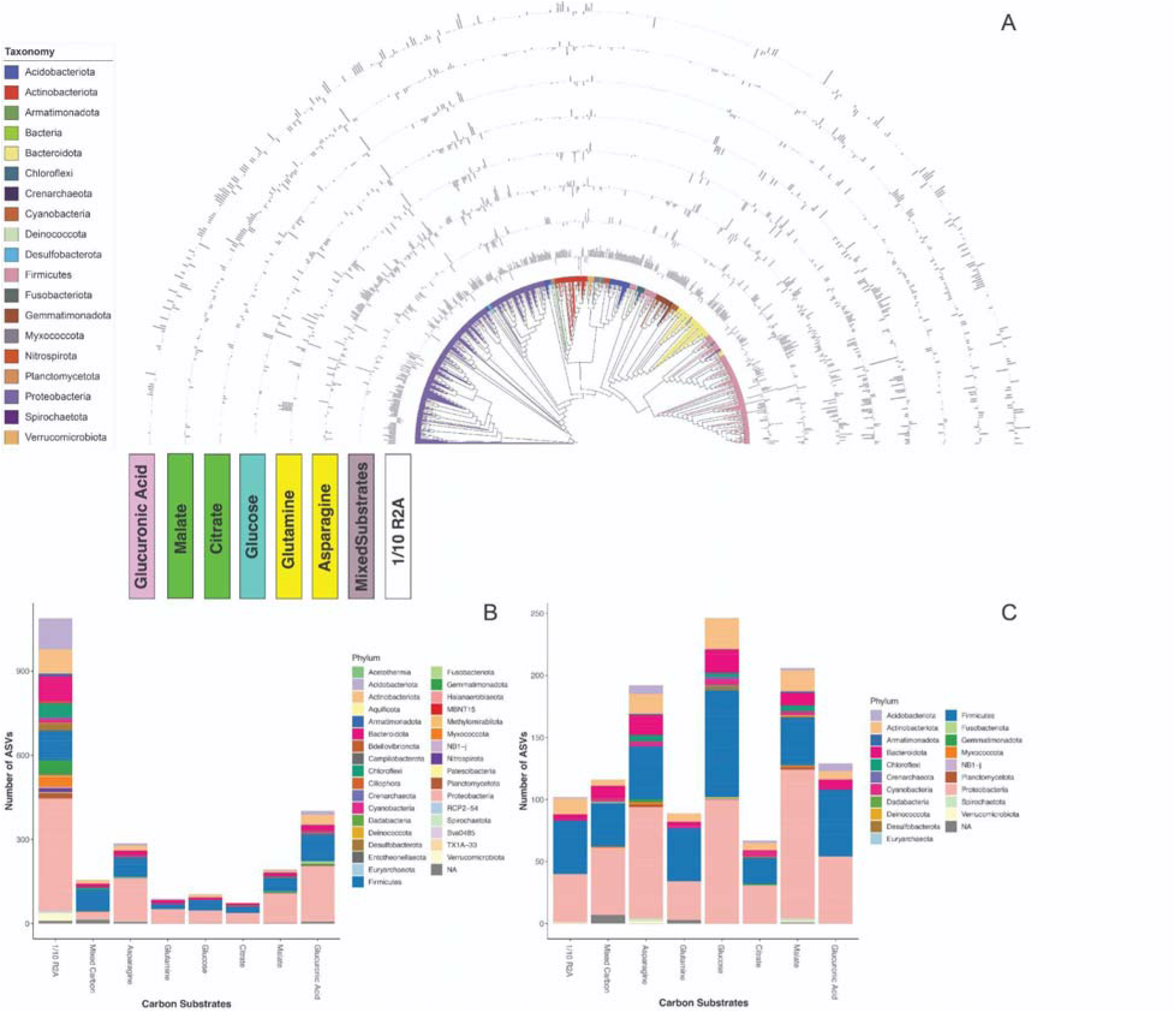
(a) Neighbor-joining tree of 530 differentially abundant ASVs significantly enriched either in 7-day or 3-day enrichments from different carbon substrates. Bar charts around the tree represent log-fold changes for each OTU. number of ASVs that are significantly enriched in (b) 7-day or (c) 3-day enrichment from different carbon substrates, different colors represent different phyla.

Additionally, we used network analyses to identify the key taxa of microbes with different growth rates. When comparing core-microbes in different time intervals, we obtained 555 distinct core ASVs for 3-day intervals and 254 distinct core ASVs for 7-day intervals. We used Jaccard indices to measure the similarity between the sets of most central nodes and hub nodes in the two centers (Supplementary Table S4). There were no differences between the most central nodes of the two networks, which would be indicated by a small probability P (LJ≤jL). Similarly, the adjusted Rand index (0.305, *p-value* = 0) indicates a high similarity of the two clustering patterns. However, the hub taxa of the two networks were completely different. ASVs belonging to family Yersiniaceae, genus *Promicromonospora*, *Phyllobacterium*, *Pelosinus*, and *Sphingomonas* are hubs for 3-day interval transfers, while ASVs from genus *Terriglobus*, *Lactobacillus*, *Bacillus*, and *Mesorhizobium* are hubs for 7-day interval transfers (Figure 4).

**Figure 4:**
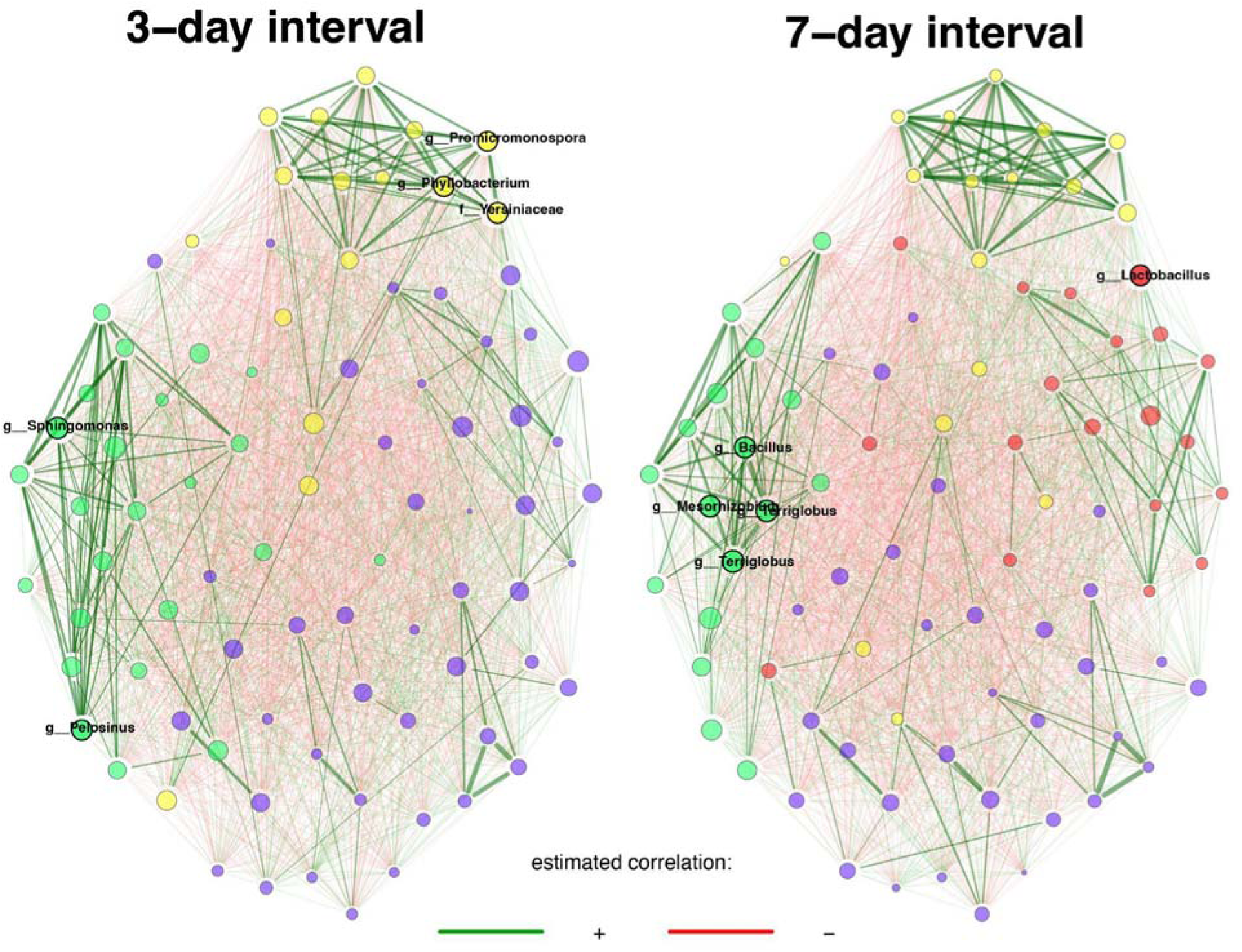
Comparison of bacterial associations between samples from 3-day and 7-day transfer intervals. Pearson correlation (>0.3) is used as an association measure, and t-test (< 0.05) is used as a sparse matrix generation method. Eigenvector centrality is used for defining hubs and scaling node sizes. Node colors represent clusters, which are determined using greedy modularity optimization. Clusters have the same color in both networks if they share at least two taxa. Green edges correspond to positive estimated associations and red edges to negative ones. The layout computed for the 3-day interval network is used in both networks. Only nodes from clusters with more than 10 nodes are considered in the plot. Hubs are highlighted with bold boundaries and corresponding taxon names.

### Evaluation of the efficacy of constructed reduced complexity consortia

Finally, we examined the efficacy of our pipeline for constructing reduced complexity consortia by performing a series of stability tests. We first compared the stability and richness of enrichment samples from different transfer generations. Most of the dominant taxa (genus > 1% in at least one sample) from different carbon sources showed no significant changes in relative abundance over time, except for samples amended with 1/10 R2A (Supplementary Figure S2). In terms of the volatility of samples from different timepoints, the mean distance between generations from all samples is 0.11. The volatility of most carbon substrates did not decrease with later transfer, except for mixed carbon substrates (Figure 5). However, allccarbon substrates except for 1/10 R2A have mean values that are lower than the global mean values of different generations (Figure 5). The species richness of most 7-day samples tended to remain stable between generations, while the species richness of 3-day samples varies more between generations (Supplementary Figure S5). In addition, the direction in which richness changes over generations varies between carbon substrates. The richness of citrate and glutamine has remained relatively stable or decreased across generations, while the richness of other carbon substrates has significantly increased in one of the later transfers (Gen 3, 6, or 9) (Supplementary Figure S5).

**Figure 5:**
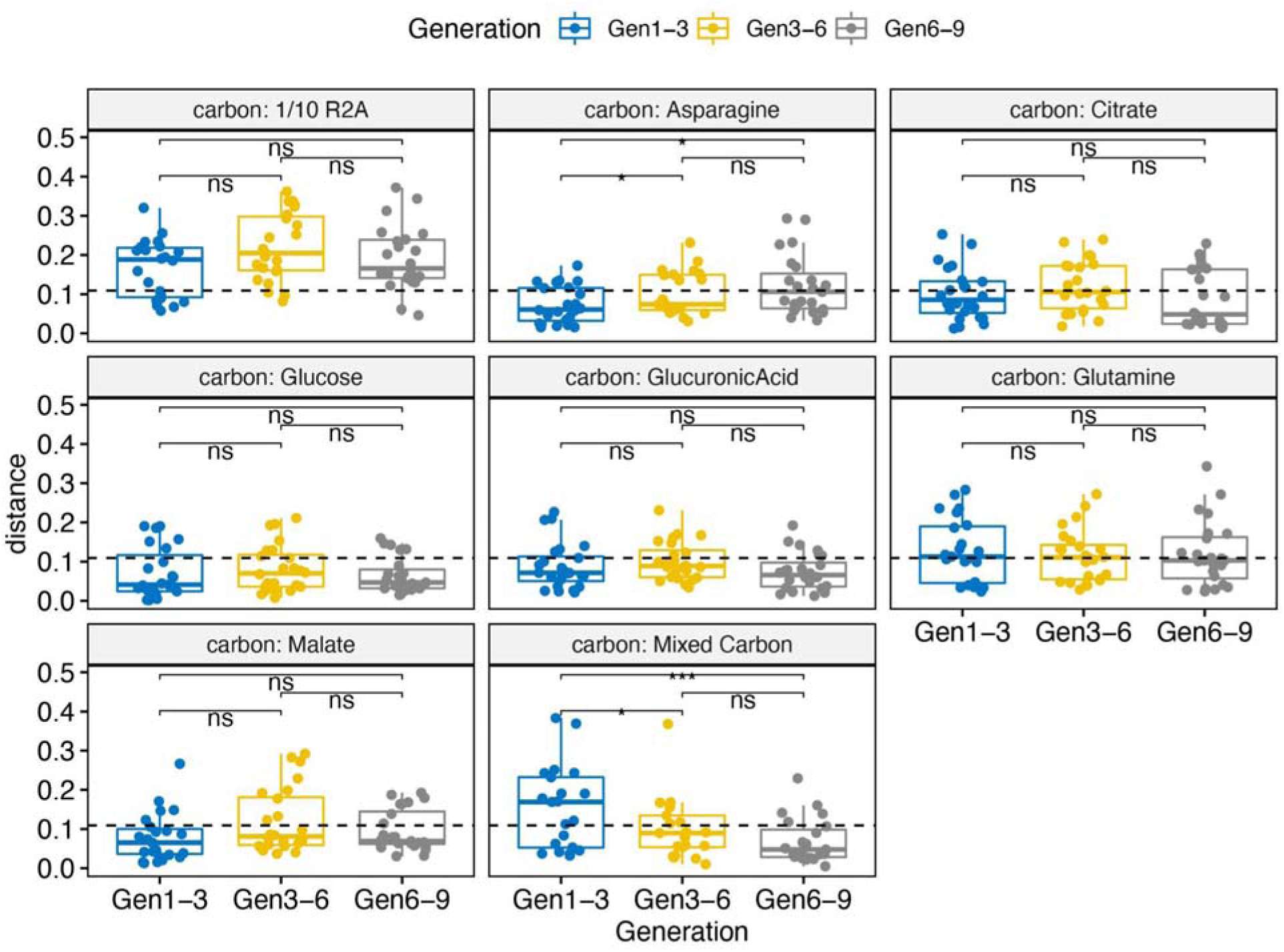
Volatility boxplot changes for samples amended with different carbon substrates comparing different generations. Gen 1-3 means differences between generation 1 and 3 which applies to other naming of x-axis. Significance code: ***: 0.001, **: 0.01, *: 0.05, ns: not significant

We further evaluated the stability, reproducibility, and revivability of our derived reduced complexity consortia by thawing five glycerol stocks with diverse microbial composition from enrichment experiments. We serially transferred these consortia five times and compared the microbial community composition among each transfer. The dominant genera (> 0.5% in at least one sample) and their relative abundances remained consistent between transfers (Figure 6). Notably, some difficult-to-cultivate genera, such as *Cohnella* and *Terriglobus*, had low but consistent relative abundances (0.1-0.5%) across different transfers. Additionally, richness and evenness of different transfers from the starting inocula showed changes within samples (Supplementary Figure S6). PERMANOVA results demonstrated that different starting inocula played a dominant role in shaping the microbial composition (*R*^2^ = 0.733, *p* = 0.001), while different serial transfers had insignificant impacts (*R*^2^ = 0.081, *p* = 0.057). Regarding the volatility of samples from different transfers, the mean weighted unifrac distance of different transfers within samples was 0.042, and many of these (17 out of 25) had a distance < 0.05 (Supplementary Figure S7). The revived samples also show a similar microbial composition to the original subculture shown by low volatility (Supplementary Table S6).

**Figure 6:**
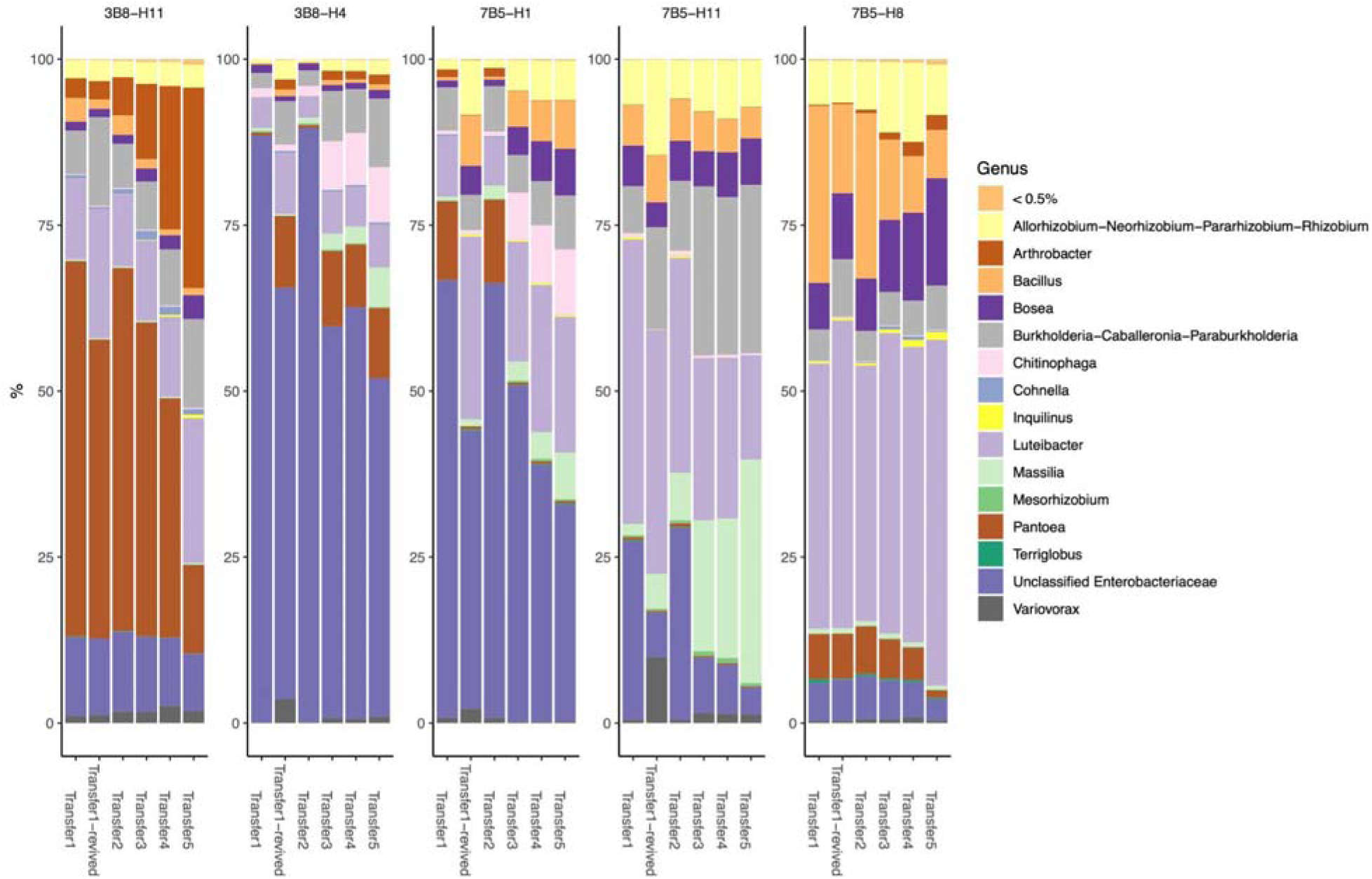
Temporal community structures of stability test of derived SynComs, reported as relative abundance of taxonomic genera (> 0.5% relative sabundance in at least one of the sample) from five different samples and five different transfers

We then employed network analysis to discern key taxa and the functional microbial communities with varying growth rates within our derived consortia. Upon comparing core microbes across different time intervals, we identified 26 unique core ASVs for both 3-day and 7-day intervals. In line with prior network analysis findings, there were no significant differences between the central nodes of the two networks, as suggested by a small probability P (J≤j) (Supplementary Table S7). The adjusted Rand index further supports this, presenting a high similarity between the two clustering patterns (0.531, *p-value* = 0). Nevertheless, variations in node shapes and edge weights for identical groups imply that the same nodes might have different interactions with others over different time intervals. For instance, the nodes for the *Cohnella* and *Chitinophaga* genera are substantially larger and have higher edge weights in 3-day intervals. Conversely, the *Terriglobus* genus node exhibits higher edge weights in 7-day intervals (Supplementary Figure S8).

## 4. Discussion

A major innovation in our pipeline is the implementation of systematic enrichments to examine how starting inocula, substrates, and time can influence the structure of microbial communities derived from enrichment experiments. Notably, the influence of time on generating synthetic communities is not well studied [45, 46]. In our study, we used the rhizosphere (including both loosely and tightly bound soil aggregates) of *Brachypodium distachyon* from various containers as the original inoculum and investigated the effects of different root exudate compounds on the overall and specific development of microbial communities. All tested parameters (original inoculum, carbon substrates, and transfer intervals) significantly impacted microbial community compositions (Figure 1). Generally, microbial communities from EcoFAB-derived inocula were distinct from those sourced from pots and tubes (Supplementary Figure S1). This could be due to a couple of factors: (1) the combination of root tip and base from different growth chambers for the initial inoculum derived from glycerol stocks of our previous study [25]; and (2) physical differences like humidity and space limitations between EcoFAB and conventional containers, which could impact root architecture [47], alter root exudate patterns, and subsequently affect the rhizosphere microbiome. Nevertheless, the original inoculum significantly impacted the richness of derived consortium in previous studies [48], and thus highlight the need to consider the impacts of inoculum when developing simplified microbial consortia.

It is important to note that other carbon substrates can also selectively enrich specific ASVs. We selected 6 single carbon substrates and mixed carbon that are dominant root exudates from *Brachypodium distachyon*, for sequencing. We also included a diluted version of commercially available undefined media 1/10 R2A for comparison. Asparagine, the dominant root exudate from *Brachypodium distachyon* [49], can significantly enrich ASVs from genera *Bradyrhizobium* and *Allorhizobium-Neorhizobium-Pararhizobium-Rhizobium* (Supplementary Table S5). Both *Bradyrhizobium* and *Allorhizobium-Neorhizobium-Pararhizobium-Rhizobium* are known genera from nodule-forming microsymbionts (collectively termed “rhizobia”) associated with plant roots, promoting plant growth [50, 51]. These results illustrate that dominant root exudates in the rhizosphere selectively enrich microbes beneficial to plant growth. Furthermore, other root exudate components were also successful in enriching ASVs associated with plant growth promoting traits. Glucose significantly enriched ASVs from the genera *Mycobacterium* and *Delftia*, both known for non-rhizobial endophytes or phosphate-solubilizing bacteria that promote plant growth [52]. Prior research shows that adding glucose can enrich the genus *Delftia* from the rhizosphere of *Cerasus sachalinensis* [53]. *Brevibacillus*, significantly enriched in glutamine-amended samples, contains strains classified as plant growth-promoting rhizobacteria (PGPR) that can inhibit pathogenic microbes and boost plant growth [54]. Interestingly, glucuronic acid significantly increased enrichment diversity and enriched the second-highest number of distinct ASVs (Figures 1, 2). Although the general impact of glucuronic acids on the rhizosphere is relatively unexplored, recent studies indicate that glucuronic acid positively correlates with soil bacteria and is considered a key metabolite [55]. Therefore, our findings emphasize the importance of using relevant carbon substrates when attempting to enrich rhizospheric microbial consortia for varied phylogenies and functions.

Moreover, choice of carbon substrates can distinctly influence fast- and slow-growing microbes. While the rhizosphere microbiome is typically dominated by fast-growing copiotrophs [56], slow-growing oligotrophs still play crucial roles including protecting against plant diseases [57] and combating environmental stress [58]. Many slow-growing microbes are less abundant in the rhizosphere, making direct isolation and characterization challenging without intelligent enrichment efforts [59]. Our results show that the use of different carbon substrates yields similar enrichment potentials for fast-growing microbes (Figure 3B). This could be attributed to the fact that the original inocula originating from immature roots, are often dominated by fast-growing microbes [60, 61]. However, distinct differences are observed in the enrichment potentials of various carbon substrates for slow-growing microbes (Figure 3C). Notably, 1/10 R2A substantially enriched slow-growing microbes, including the phyla Acidobacteria and Verrucomicrobia, which are often recognized as difficult to enrich microbes [62, 63] but with high potential for promoting plant growth [64, 65]. These findings underscore the importance of employing multiple carbon substrates, particularly complex carbon sources, to enrich rare but crucial slow-growing microbes from the rhizosphere.

We further explored microbial interactions of our reduced complexity consortia using network analysis, focusing on key taxa that significantly influence slow- and fast-growing microbes. Network pattern analysis allows us to determine community-level dynamics within intricate webs of species interactions and associations [66, 67], as well as select core taxa for constructing microbial consortia [68]. We discovered that while the two networks of different time intervals (3-day and 7-day) are overall similar, the hub taxa are entirely different (Figure 4). The hubs, identified using the “NetCoMi” R package, are crucial nodes (or keystone taxa) that hold central positions based on criteria such as the highest degree centrality [69]; highest degree, betweenness and closeness centrality at the same time [70]; or highest eigenvector centrality [71]. Consequently, hubs can be considered ecologically important microbes responsible for shaping the microbial community structure and dynamics [70]. The stark differences between hubs for the two networks suggest distinct keystone taxa among slow- and fast-growers. Notably, eight keystone taxa from the hubs belong to genera known to exhibit plant growth-promoting (PGP) traits [64, 72–78]. The strong association between keystone taxa and PGP traits indicate that PGP bacteria significantly influence the rhizosphere microbiome’s dynamics [79] and emphasizes the importance of selecting them when constructing rhizosphere microbial consortia. In addition, the network analysis demonstrated consistent grouping trends over different time intervals of transfer. Yet, it should be noted that each individual node may display varying likelihoods of interaction with other members within the same community, as shown in the Supplementary Figure S8. For instance, a node from the genus *Terriglobus*, which is known for slow growth rates [80], demonstrates more active interaction with other nodes over 7-day intervals. On the other hand, nodes from the *Cohnella* and *Chitinophaga* genera, recognized as fast-growing microbes [81, 82], exhibited increased interactions with other nodes over 3-day intervals. These observations underscore the importance of longer transfer times when attempting to construct consortia, to ensure the inclusion of more slow-growing microbial members.

Finally, we evaluated the feasibility of our pipeline by examining the stability, reproducibility, tractability, and revivability of our derived microbial consortia. We initially assessed the stability and richness of serially-transferred enrichments at several generations to validate our approach in building microbial consortia, as stability and tractability are two essential features for creating successful microbial consortia [17, 23, 48]. We observed relatively stable and low volatility (< 0.11, the median distance across all samples) for microbial communities between different transfers from various enrichments (Figure 5). Likewise, most enrichments demonstrated stable and low species richness (<70 observed species) across different generations (Gen 1, 3, 6, and 9), regardless of the testing variables (Supplementary Figure S5). These findings suggest that microbial communities from these enrichment experiments may have already reached stablity at the early stages of the transfer, with the exception of samples amended with 1/10 R2A. This observation is supported by the Bray-Curtis dissimilarity matrix comparing microbial communities from Gen 1 with the samples from our prior biogeography study [25] (root tip and base), as we identified significant changes in microbial compositions between the two experiments (Supplementary Figure S3). Moreover, species richness comparisons indicate that Gen 1 samples have significantly lower richness than those from the biogeography study (Supplementary Figure S4).

Consistent revival and propagation from glycerol stocks is a highly desired trait for microbial consortia to enable study across multiple labs and timescales. We tested our microbial consortia preserved in glycerol stocks to further assess their reproducibility, stability, and revivability. Generally, for multiple samples, our derived microbial consortia exhibited consistent genera (in respect to both presence/absence and relative abundances, Figure 6), richness, and evenness across different transfers (Supplementary Figure S6) and have tractable numbers (18 to 44) of observed OTUs, demonstrating good reproducibility. The low values from the volatility tests between transfers (Supplementary Figure S7) also support the stability of the derived microbial consortia. We also verified the revivability of our derived consortia, showing that the communities revived from glycerol stocks have a similar microbial composition to the original culture (Supplementary Table S6). In conclusion, our pipeline can efficiently establish stable, revivable, reproducible, and tractable consortia from the source rhizosphere microbiome.

## 5. Conclusion

In this study, we implemented a comprehensive and systematic approach using high-throughput enrichment techniques to derive microbial consortia from the rhizosphere of B. distachyon. We evaluated various factors such as root exudate substrates, initial inoculum, and transfer duration. Our enrichment experiments revealed that the original inoculum and carbon substrates are the primary variables influencing microbial communities. The contrasting enrichment outcomes between communities sourced from plants grown in EcoFABs compared to conventional containers underscore the importance of considering the initial inoculum when constructing microbial consortia. Our enrichment findings also demonstrate that diverse carbon sources can significantly enhance the abundance of specific PGPR taxa and microorganisms with different growth rates. This emphasizes the need for using multiple carbon sources during enrichments to improve the phylogenetic diversity of microbial consortia. Furthermore, we discovered that although the keystone taxa among fast- and slow-growing microbes differ, many keystone taxa exhibit close relationships with PGPR taxa, suggesting their central roles in shaping rhizosphere microbial structures. Our enrichment methods also demonstrated a significant reduction in complexity from the original inoculum, achieving a relatively stable community after only a few transfers. Stability tests using glycerol stocks preserved from our pipeline further confirmed the reproducibility, stability, and revivability of our derived microbial consortia. In conclusion, our high-throughput enrichment methods efficiently enriched key taxa essential to the rhizosphere microbiome, achieving stable and reduced complexity consortia. This provides valuable insights for selecting criteria to construct effective rhizosphere microbial consortia. Our findings have broader implications for the field, enabling the development of tailored microbial consortia for various agricultural and environmental applications, potentially improving crop productivity, disease resistance, and environmental sustainability.

## 6. Acknowledgements

This research work was funded through the Microbial Community Analysis and Functional Evaluation in Soils (m-CAFEs) Science Focus Area Program at Lawrence Berkeley National Laboratory funded by the U.S. Department of Energy, Office of Science, Office of Biological & Environmental Research Awards DE-AC02-05CH11231. Authors would like to thank Gracielle Ria Malana and Mariam Alsaid for laboratory assistance.

