## Supplementary Figures for "Decoding Root Biogeography: Building Reduced Complexity Functional Rhizosphere Microbial Consortia"

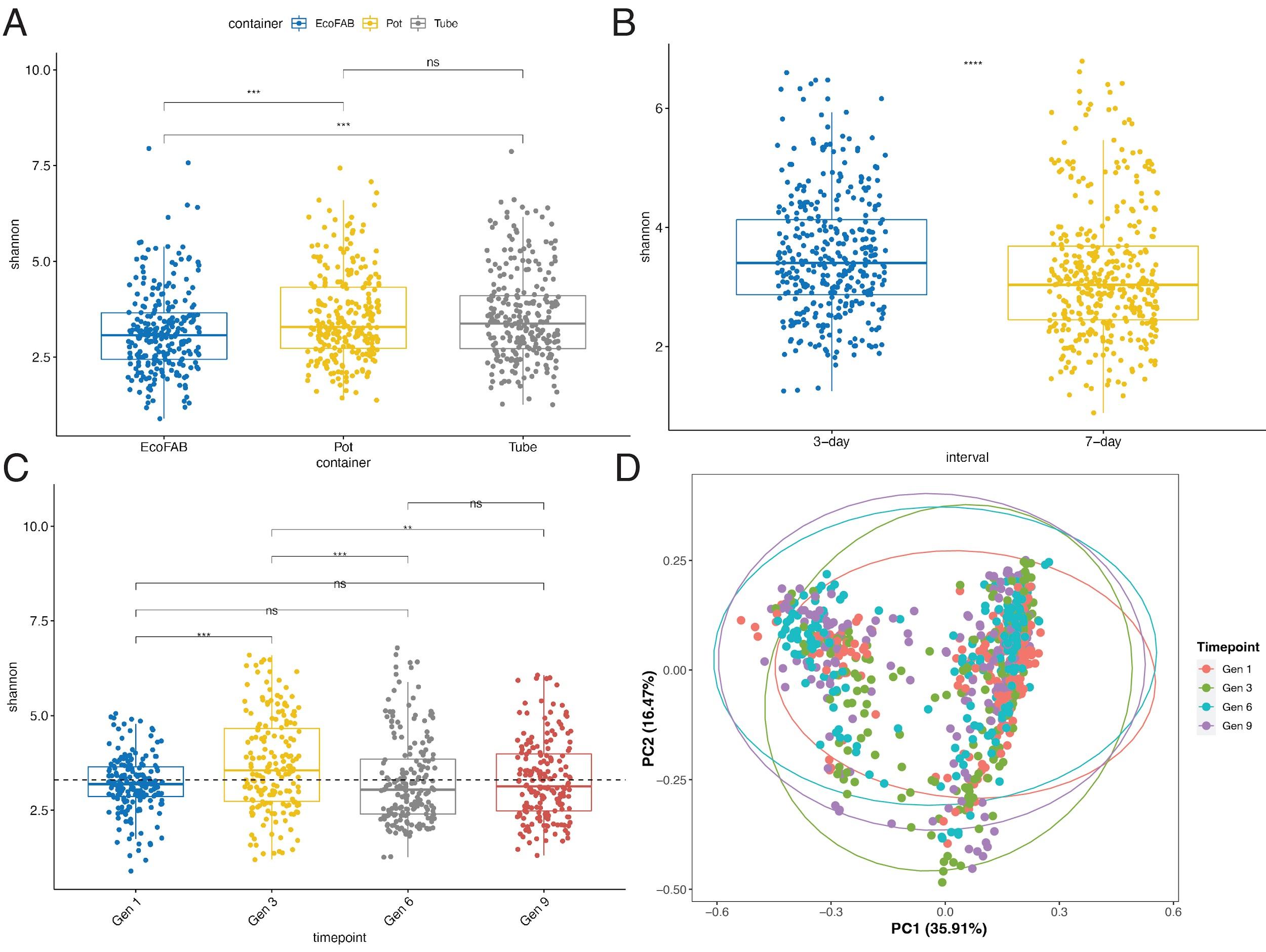

Supplementary Figure S1: Shannon Diversity comparing (A) original inoculum, (B) time intervals, (C) sampling generations , and (D) PCoA plot for sampling generations.

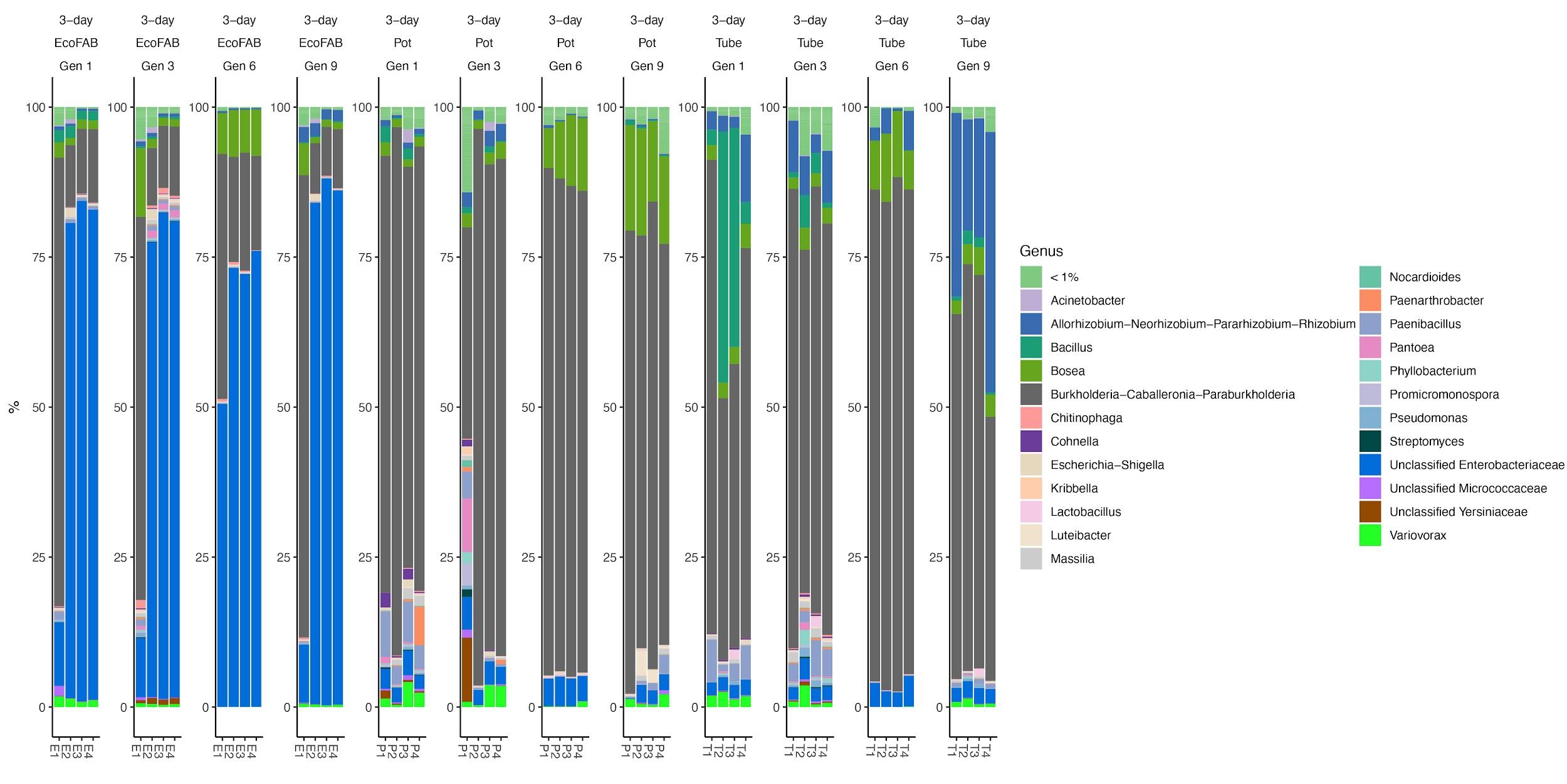

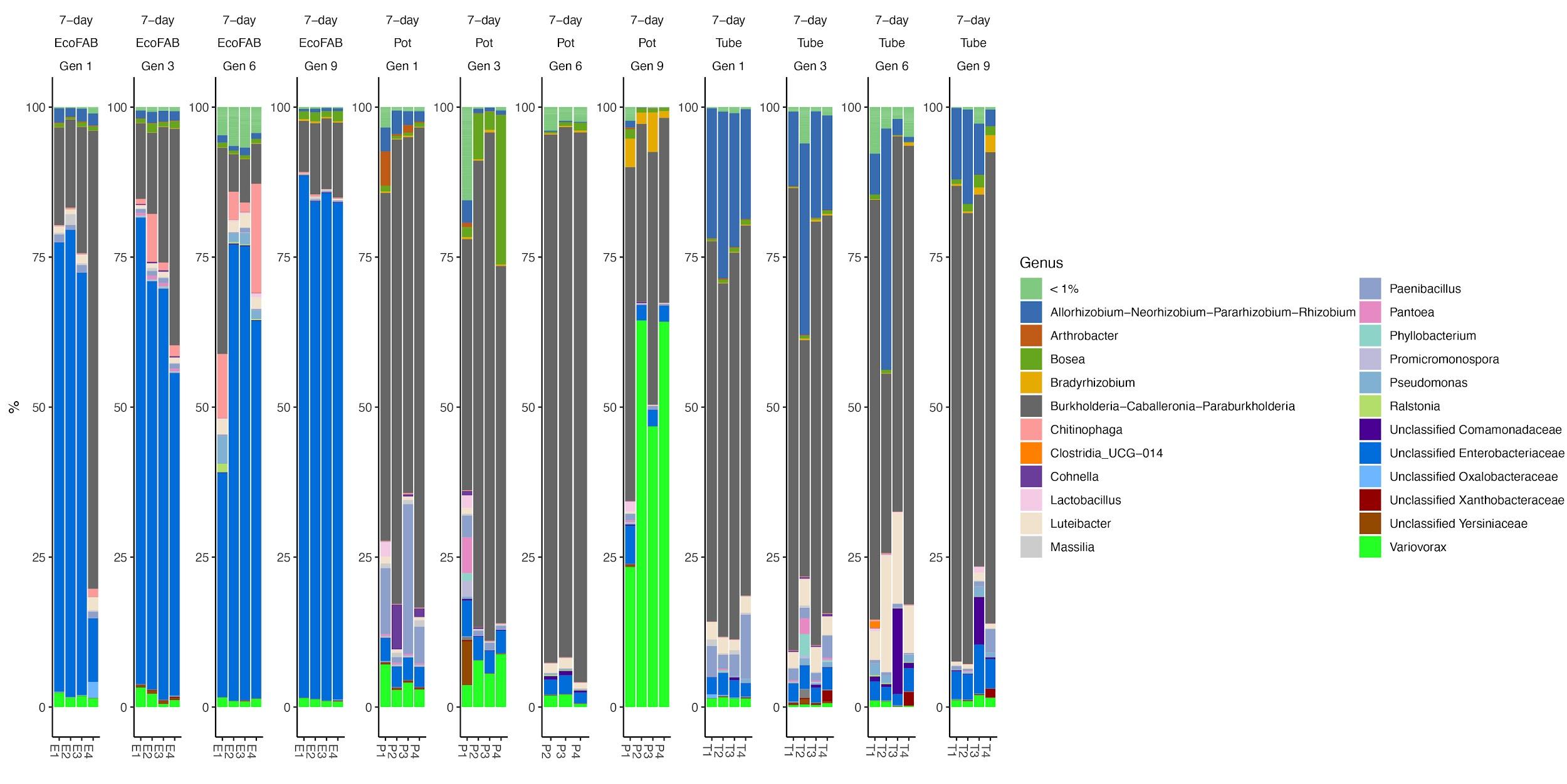

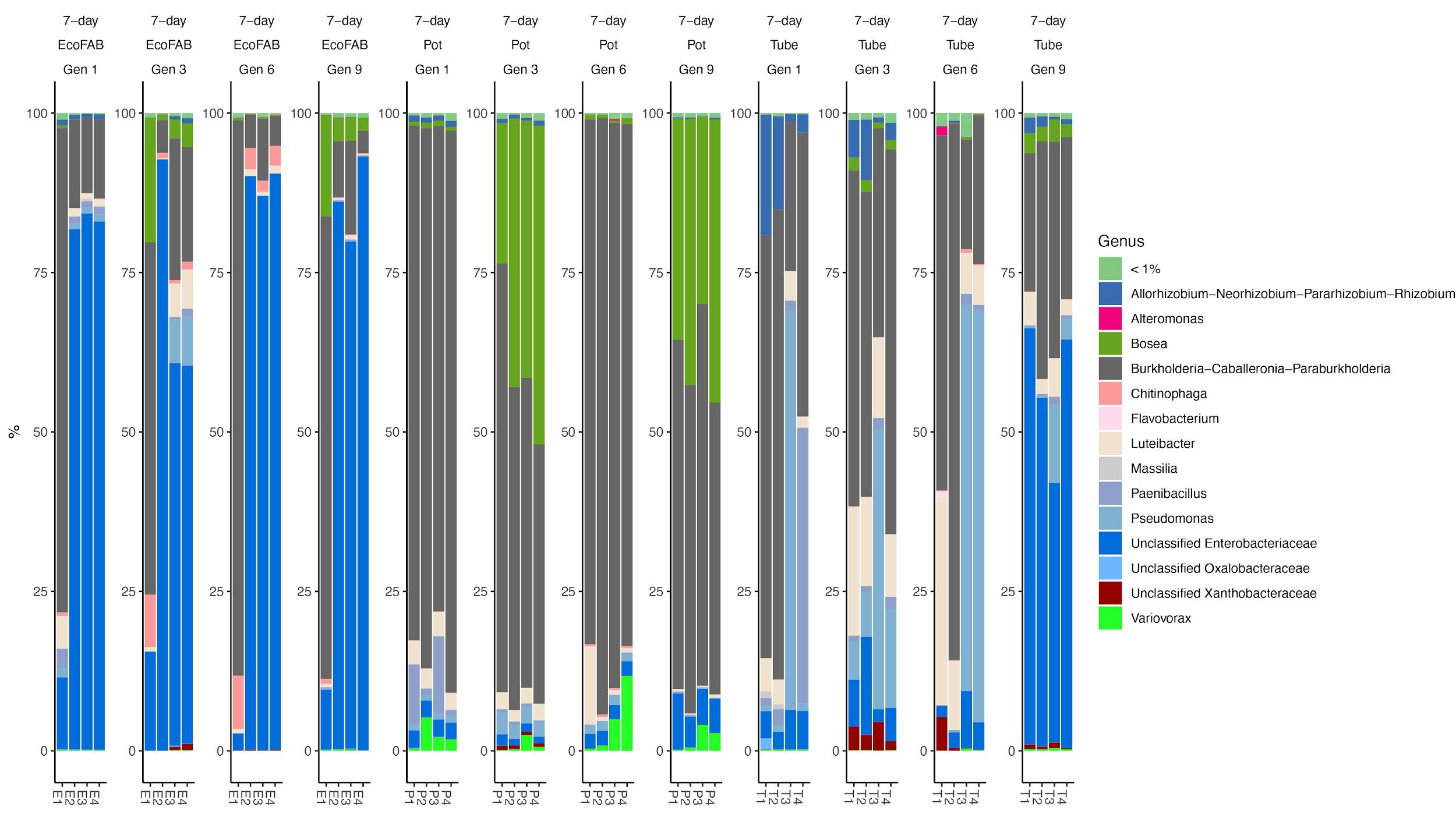

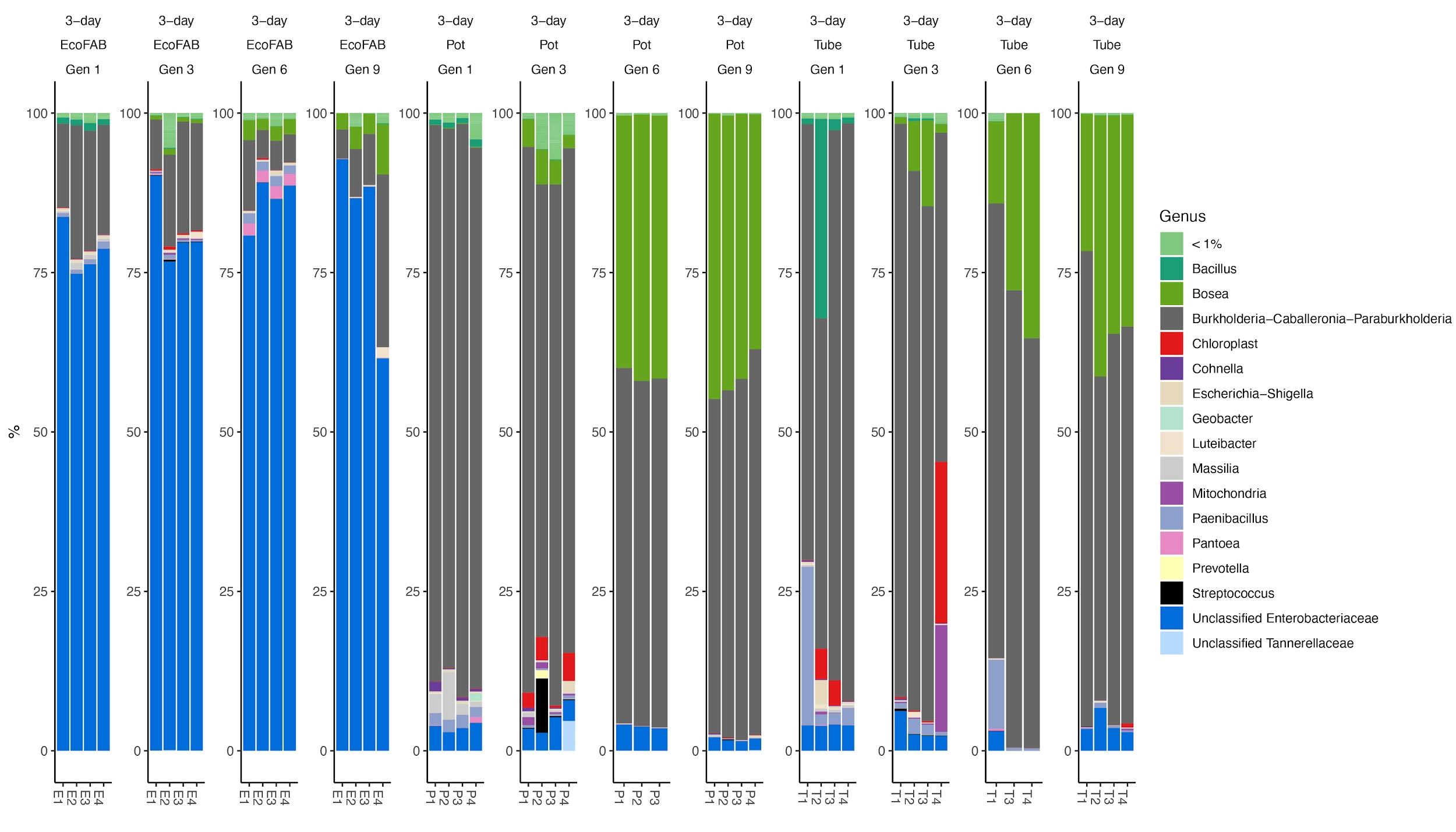

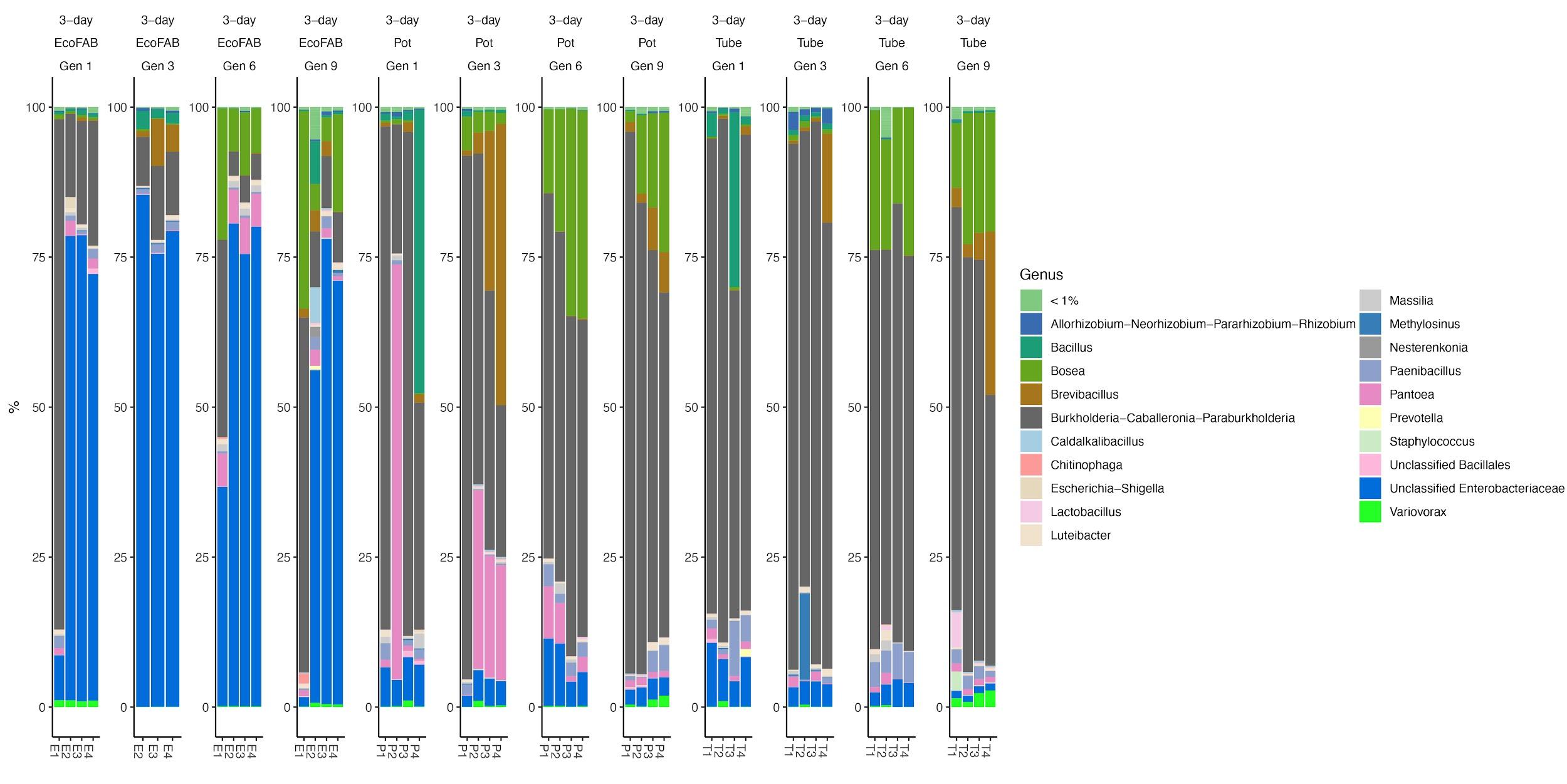

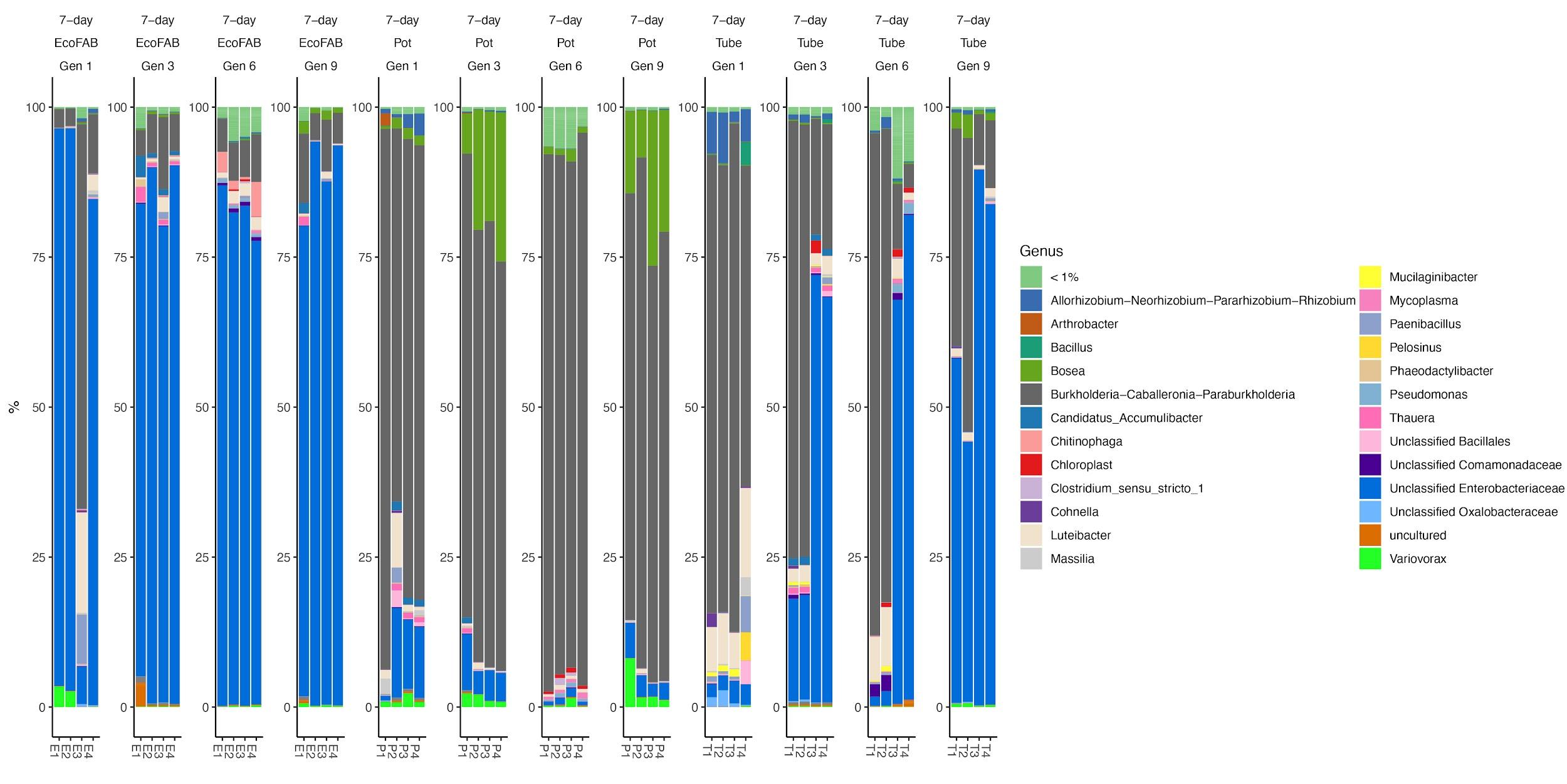

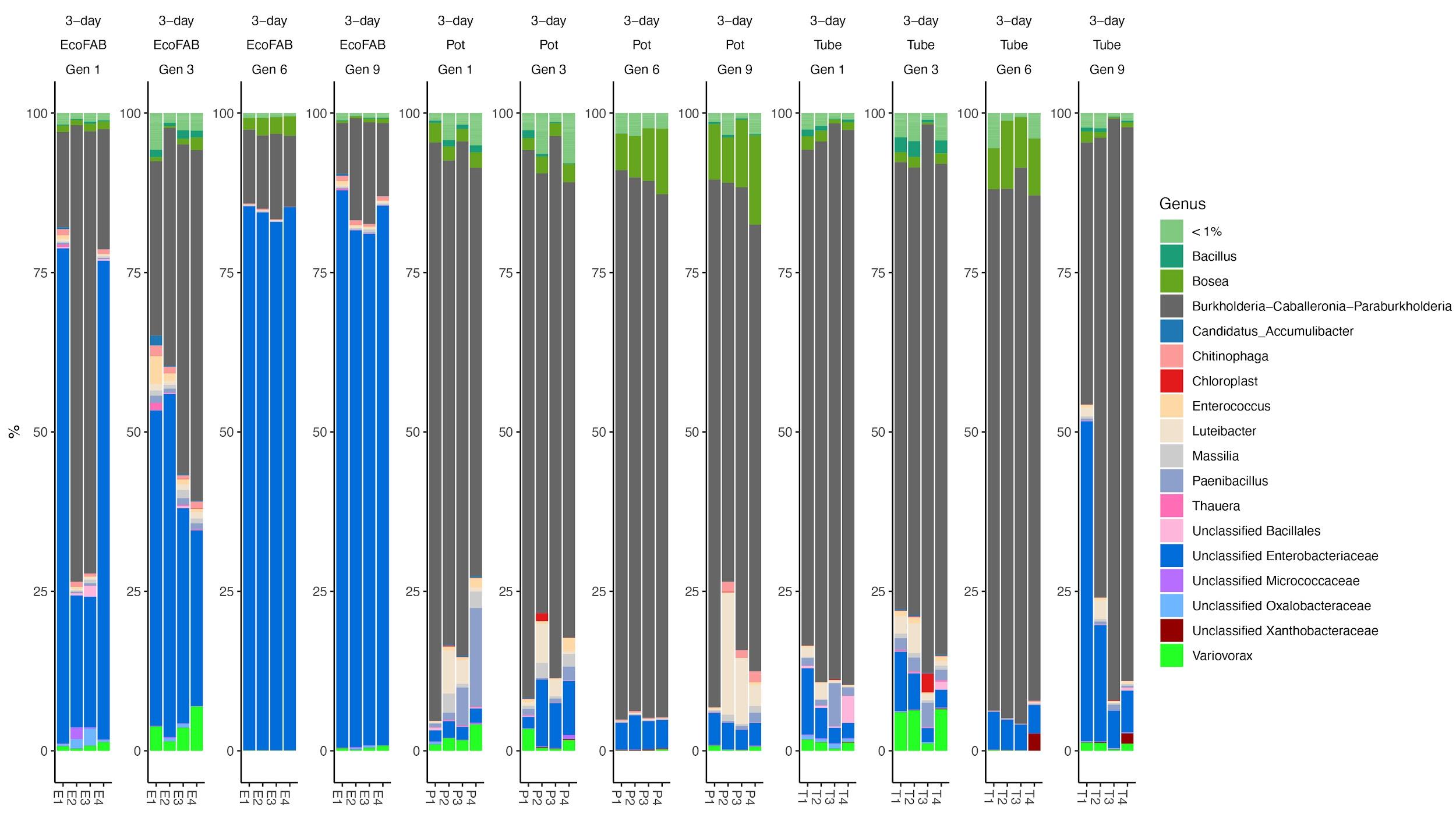

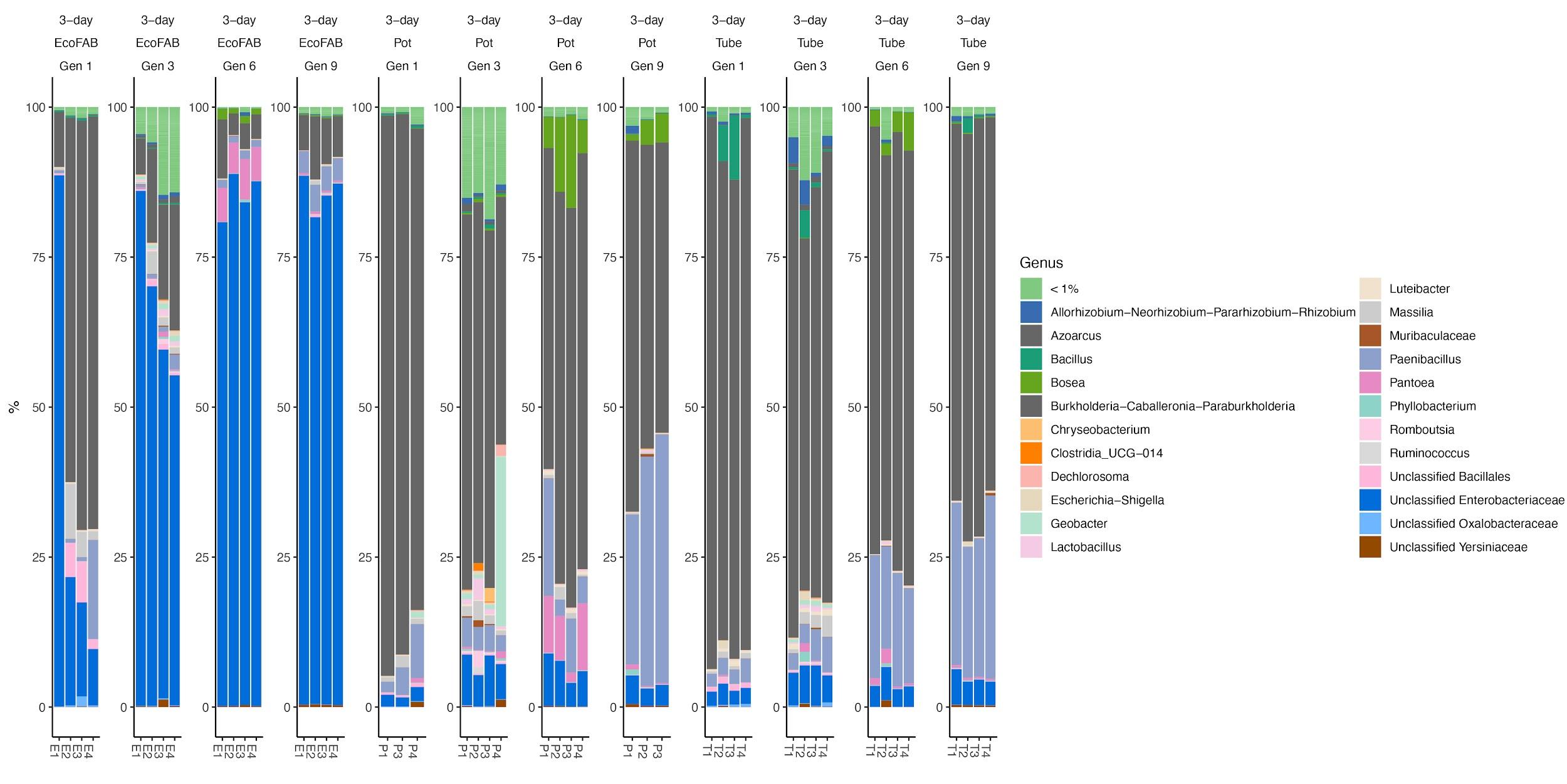

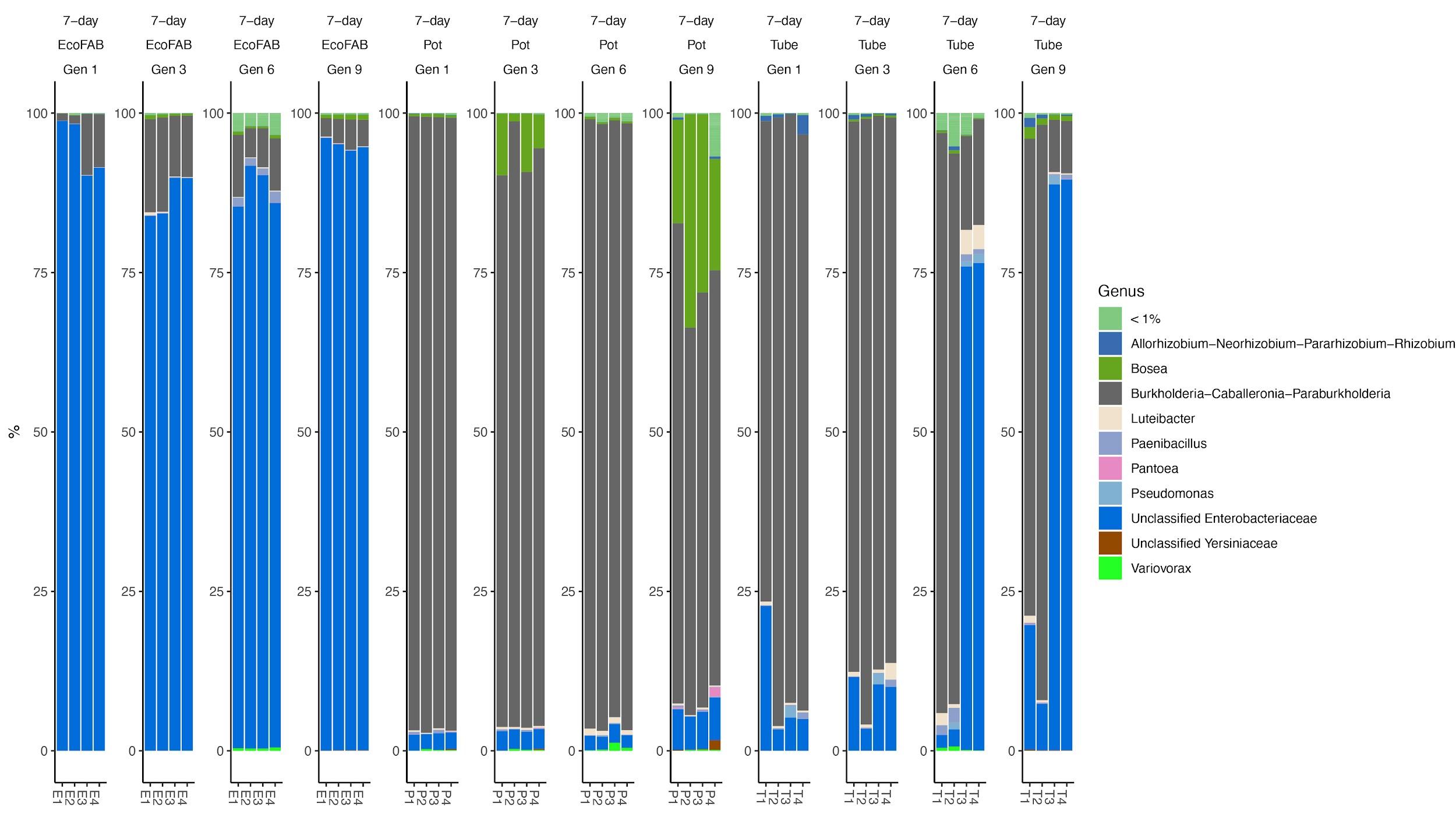

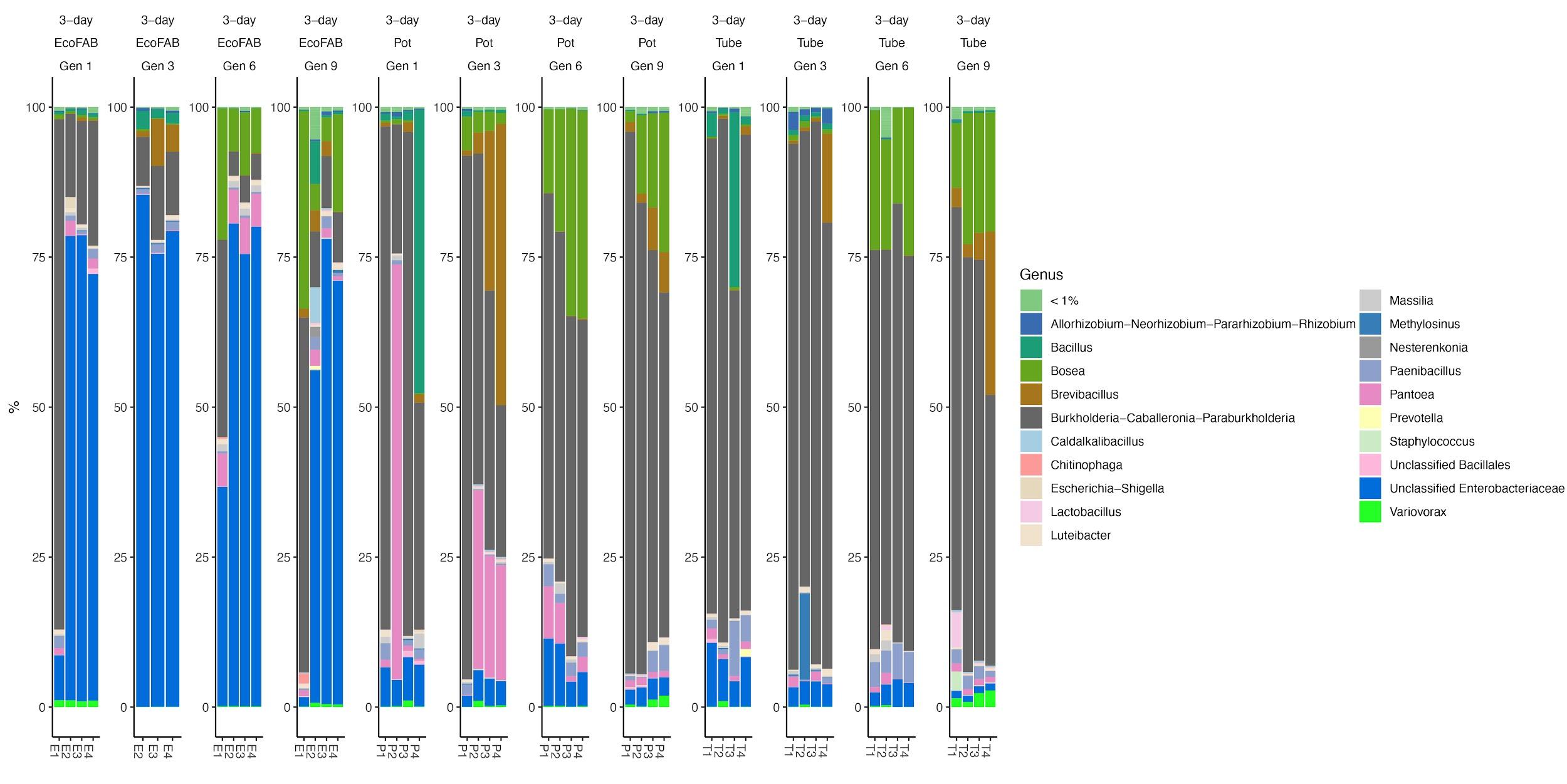

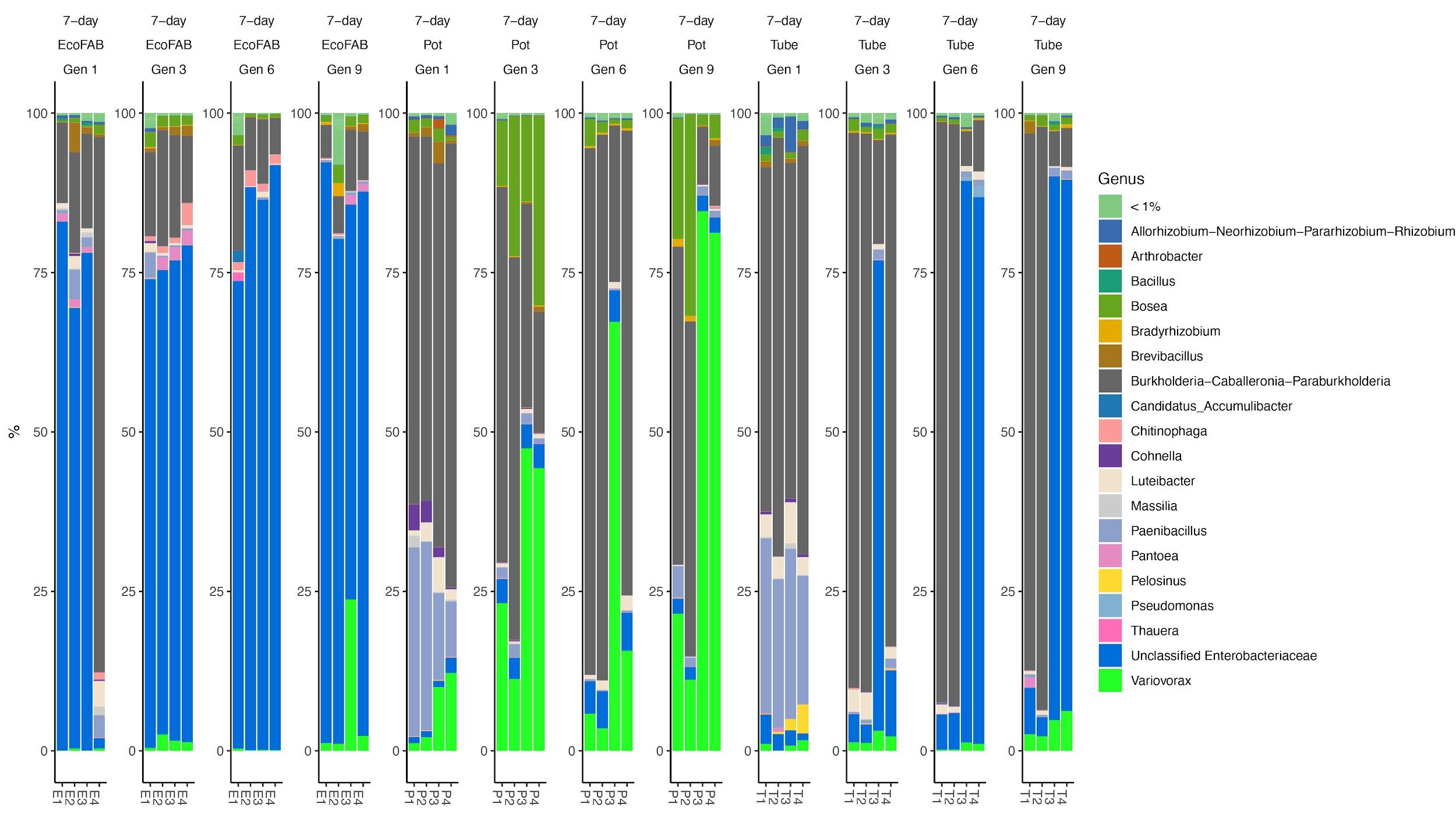

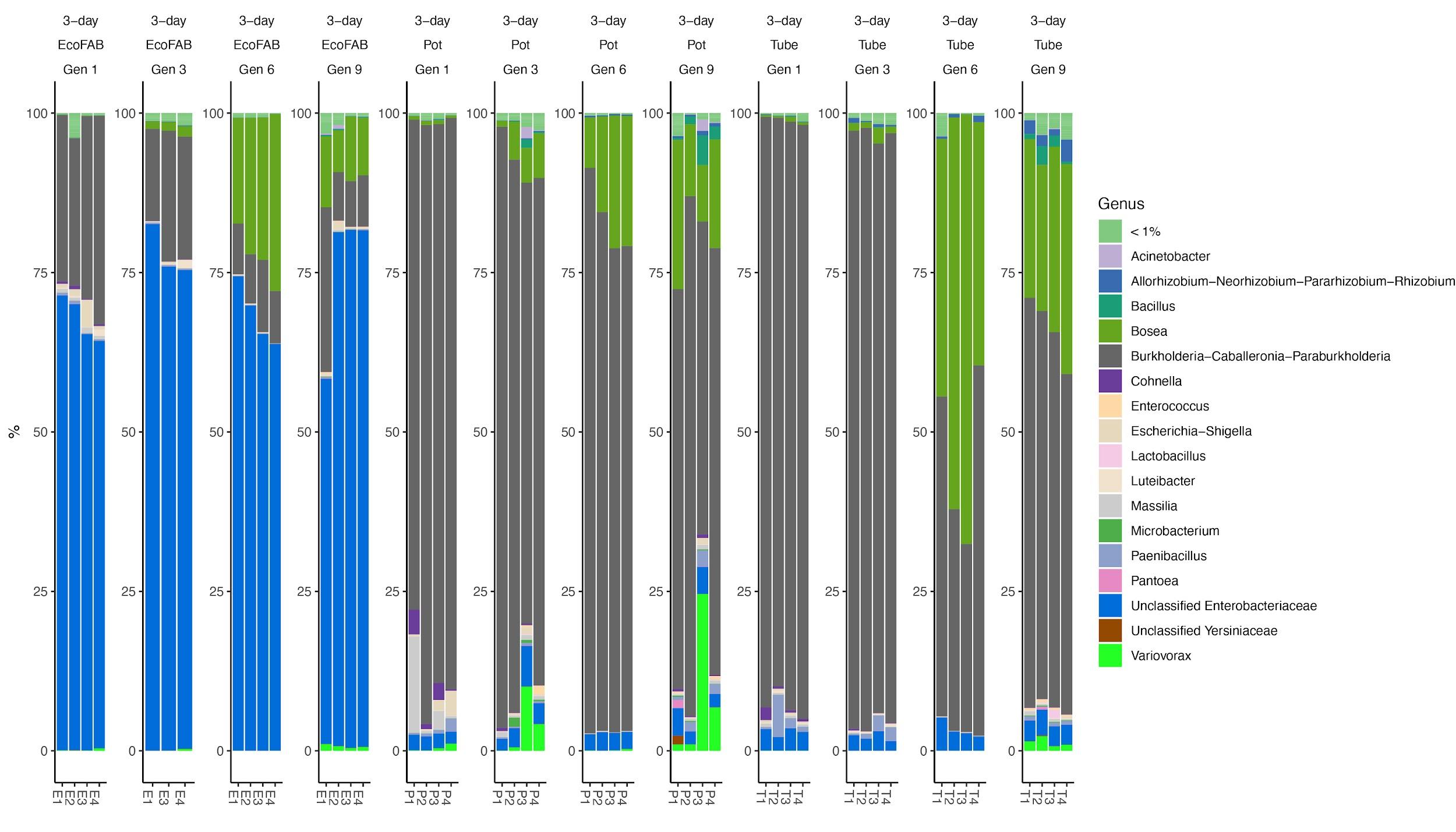

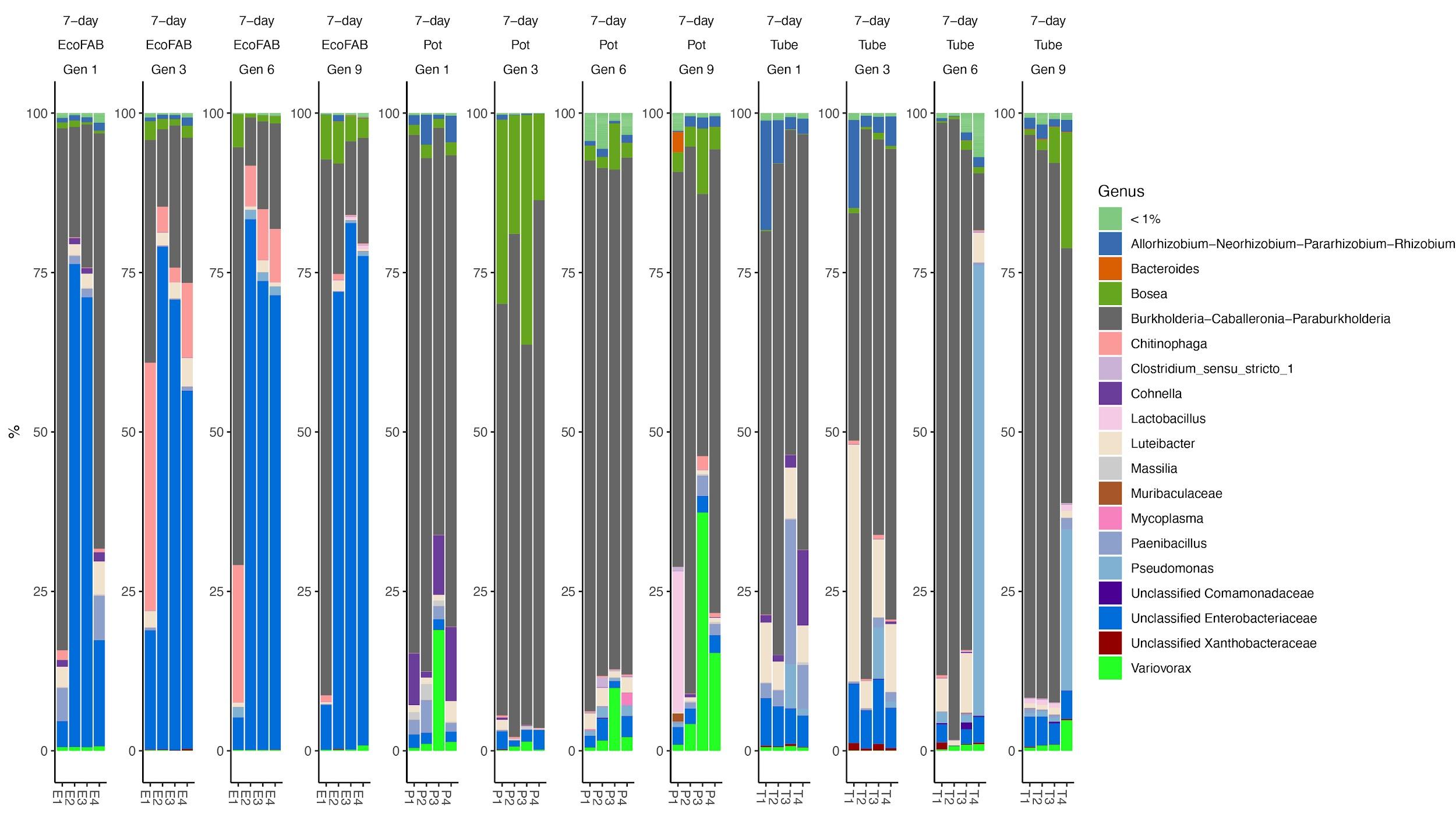

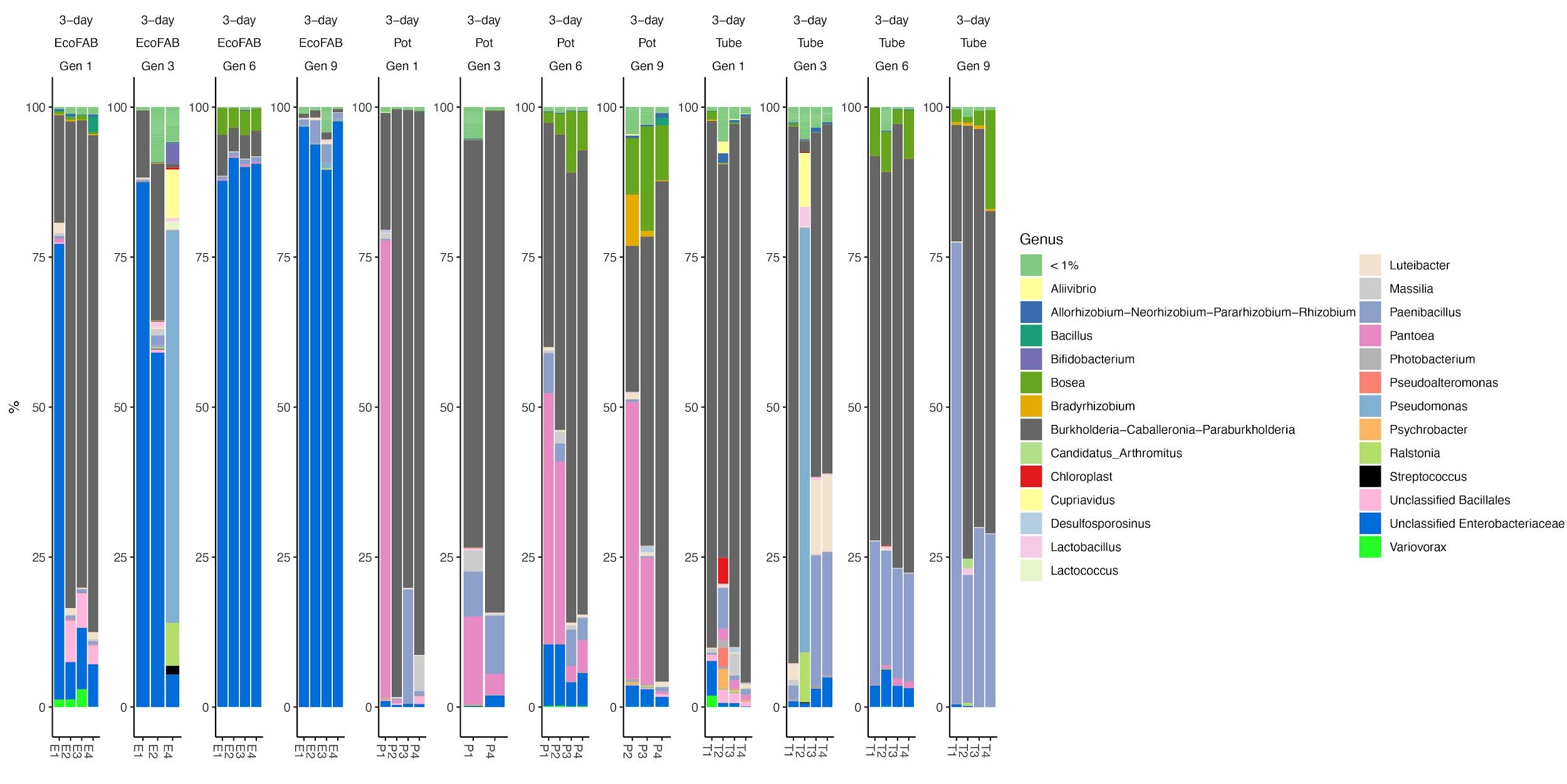

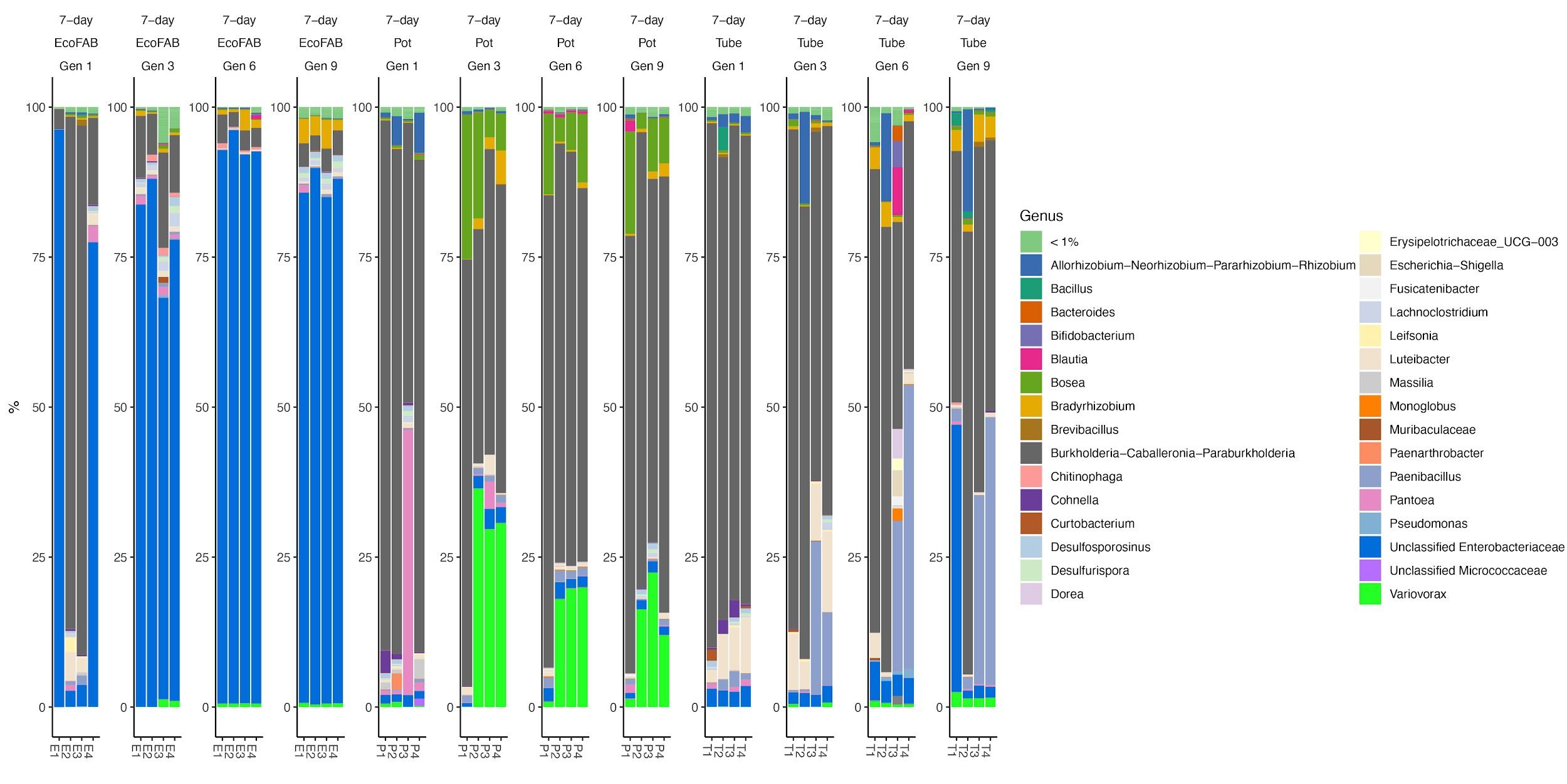

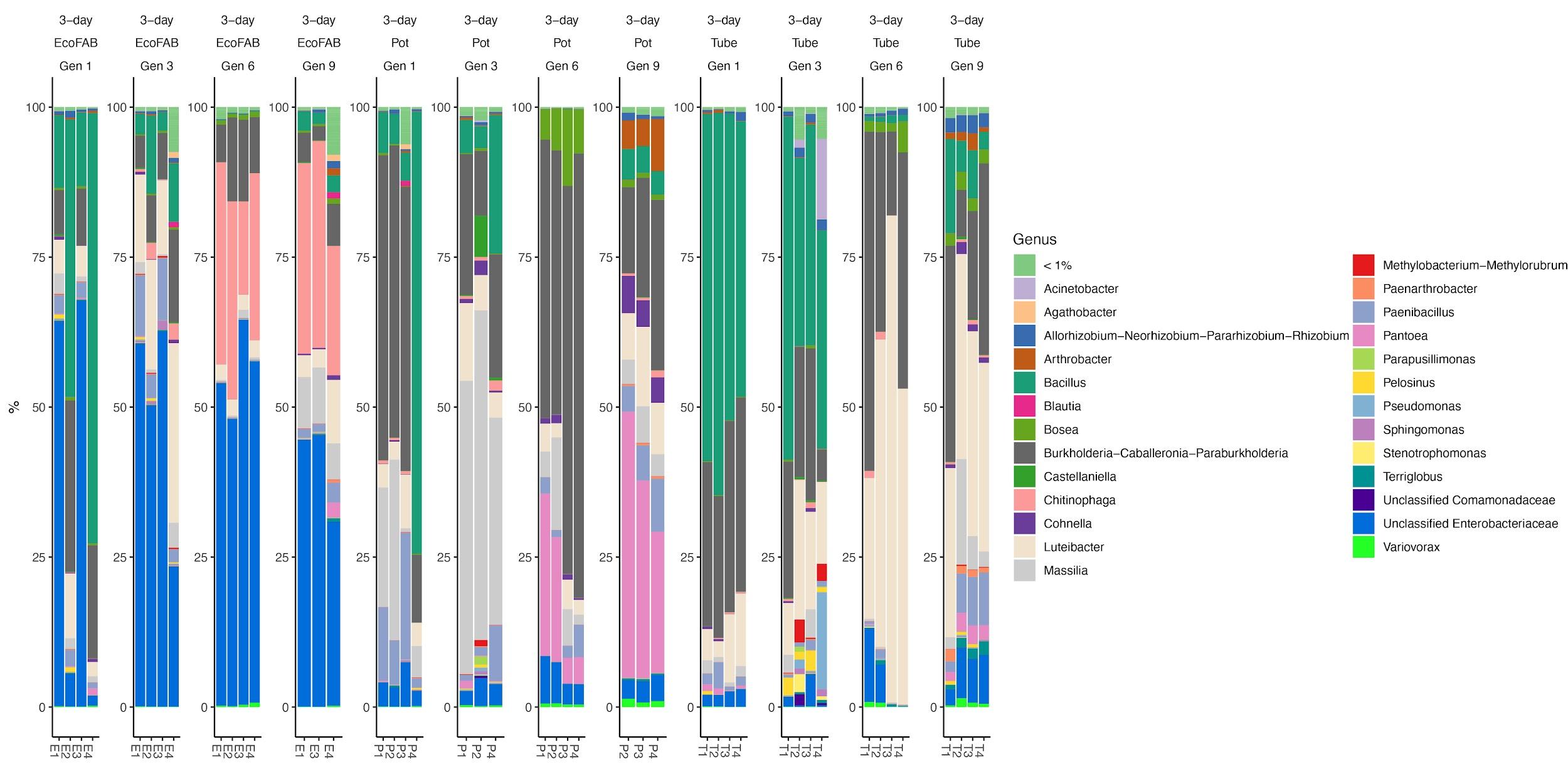

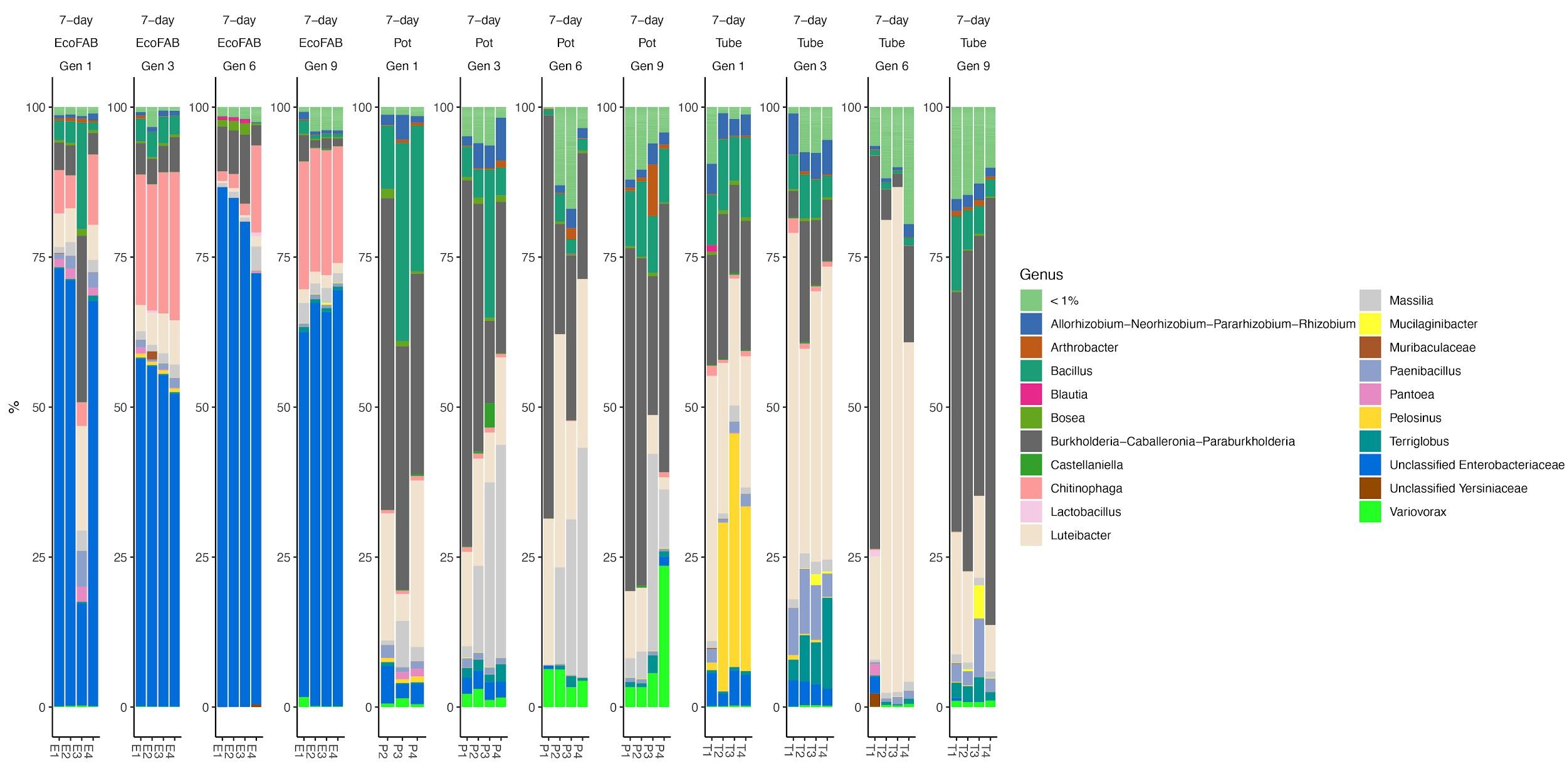

Supplementary Figure S2: Temporal community structures of different C-amended enrichments reported as relative abundance of taxonomic genera (> 1% relative abundance in at least one of the sample) from three different original inocula (EcoFAB, pot, tube), different generations and different time intervals.

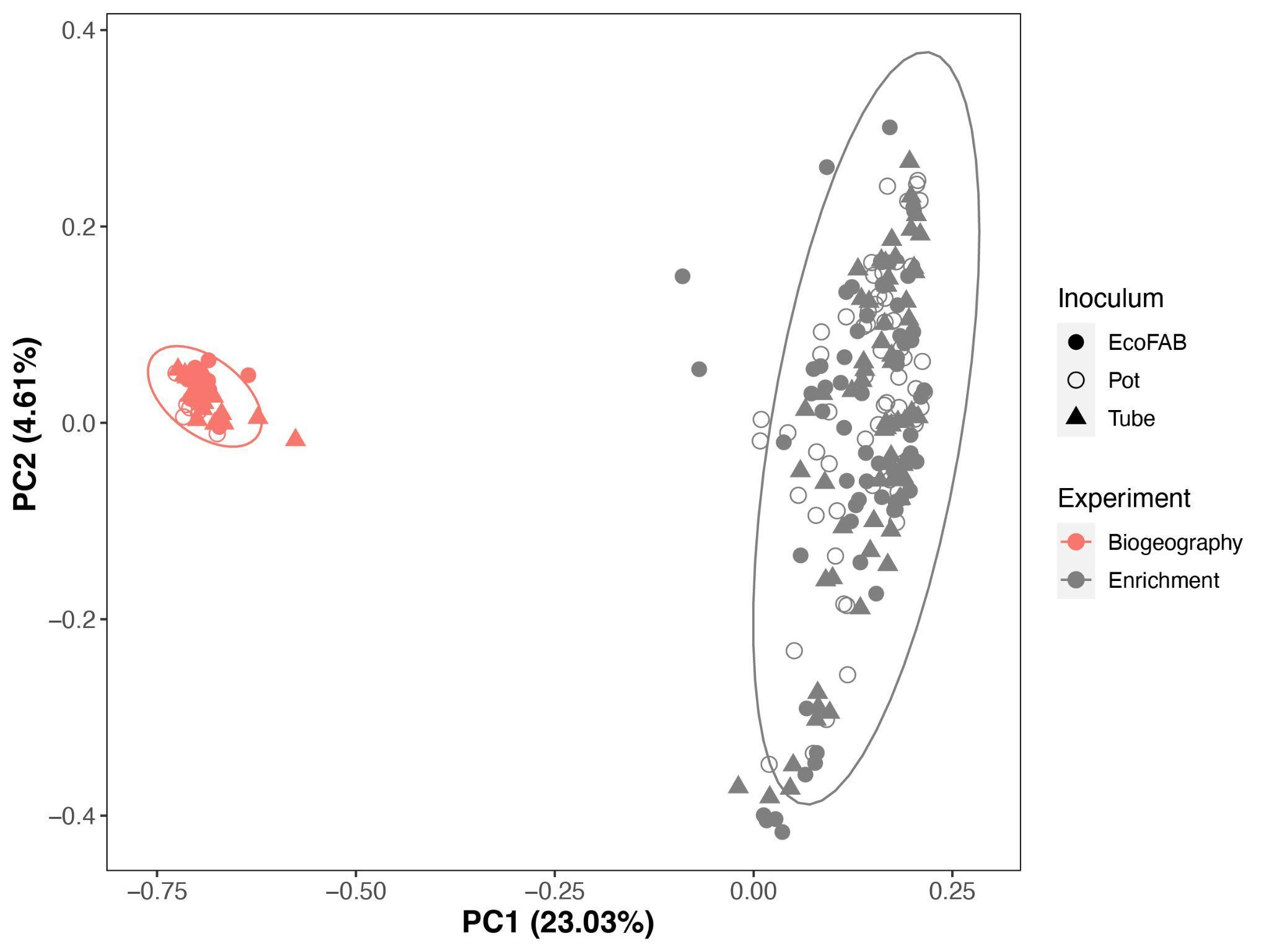

Supplementary Figure S3: PCoA plot showing the comparisons of ASVs from biogeography and enrichment experiments. The root tip and base samples from biogeography experiments and the Generation 1 samples from enrichment experiments are included in this plot.

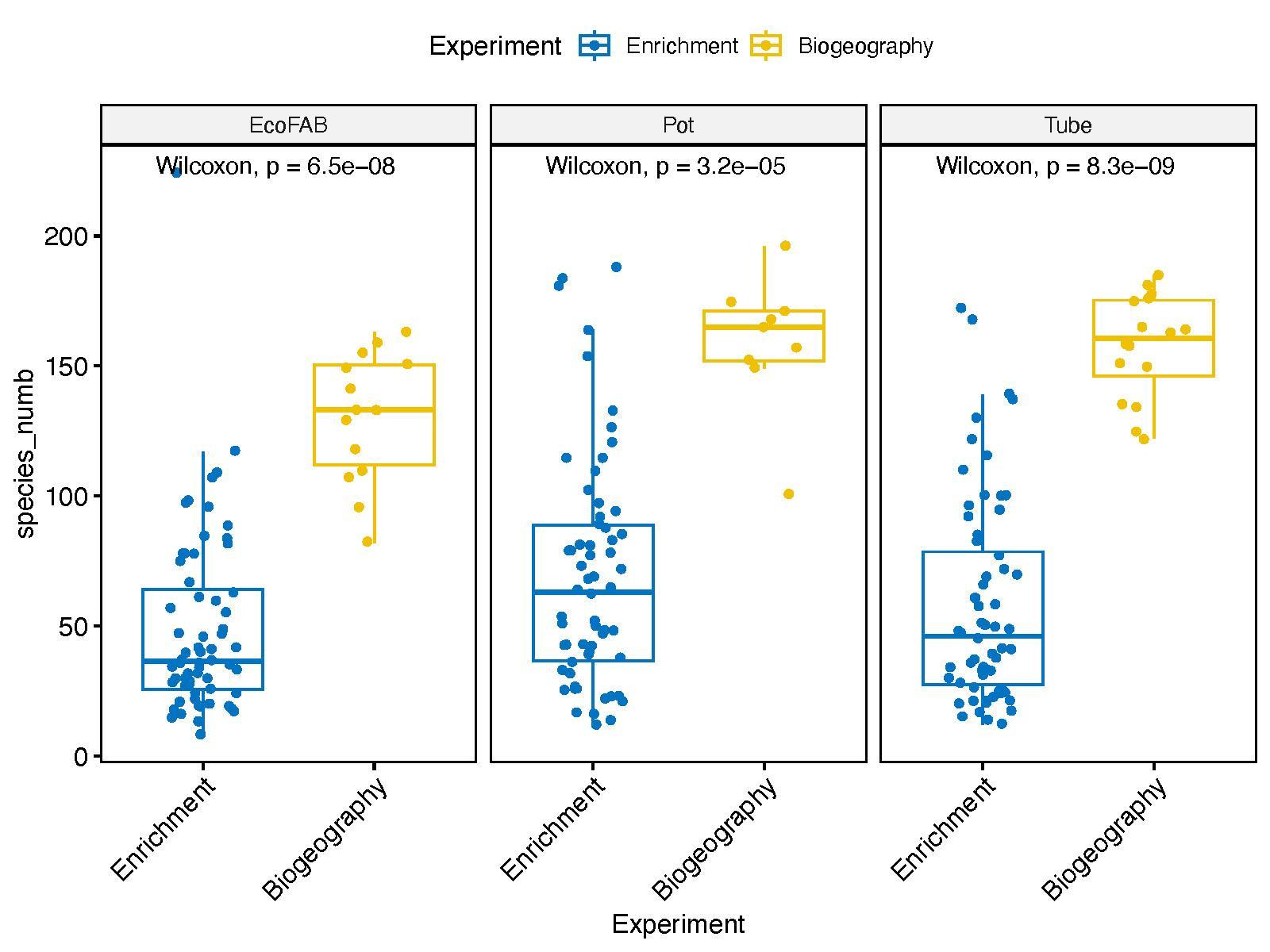

Supplementary Figure S4: Boxplot comparisons of species numbers (agglomerate to genus level) from biogeography and enrichment experiments. Wilcoxon-test p-values are included in the figure. The root tip and base samples from biogeography experiments and the Generation 1 samples from enrichment experiments are included in this plot.

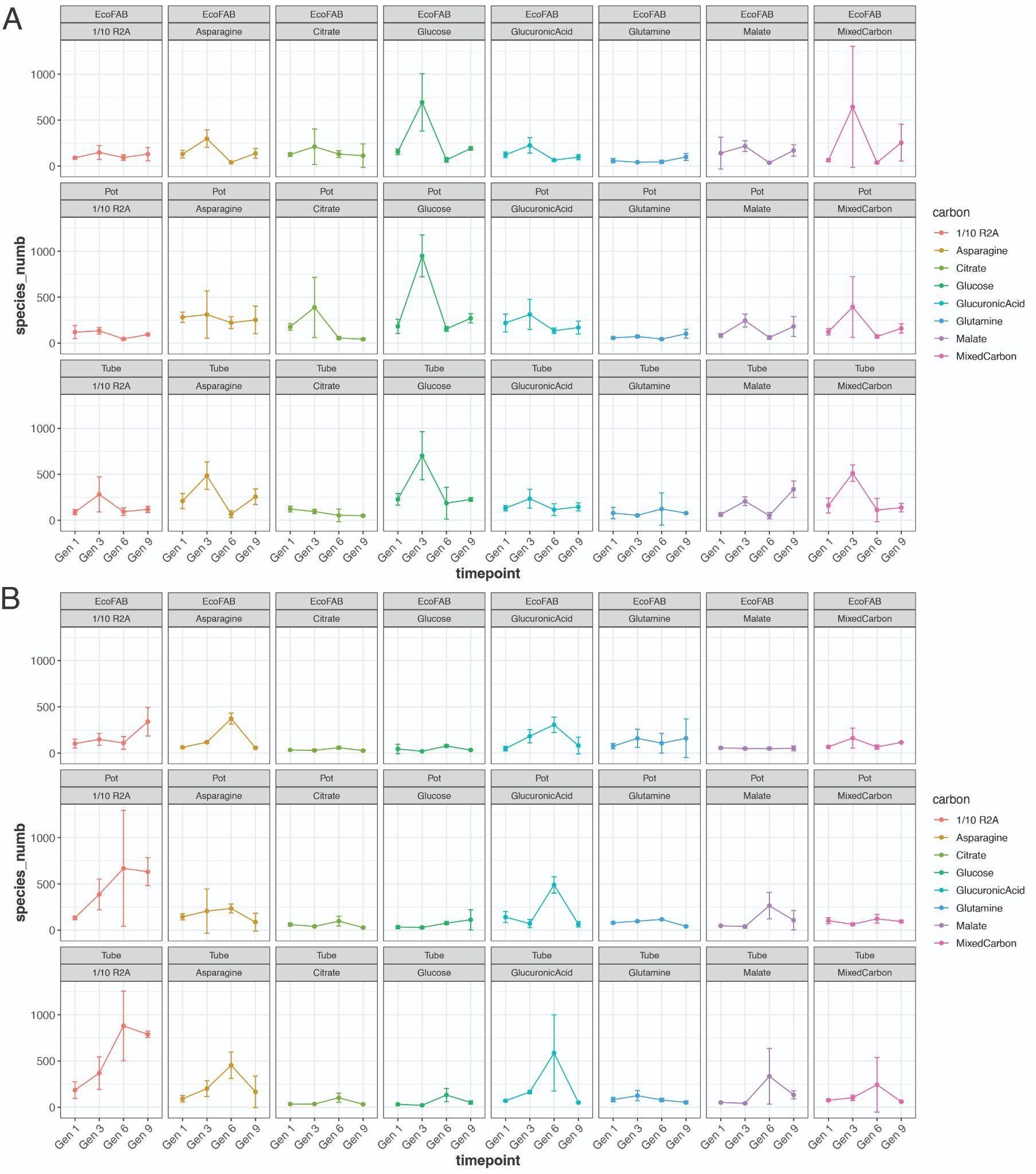

Supplementary Figure S5: Species richness boxplots for samples with different original inoculum and amended with different carbon substrates comparing different generations. Species richness numbers are depicted by dashed lines from all samples using the same original inoculum.

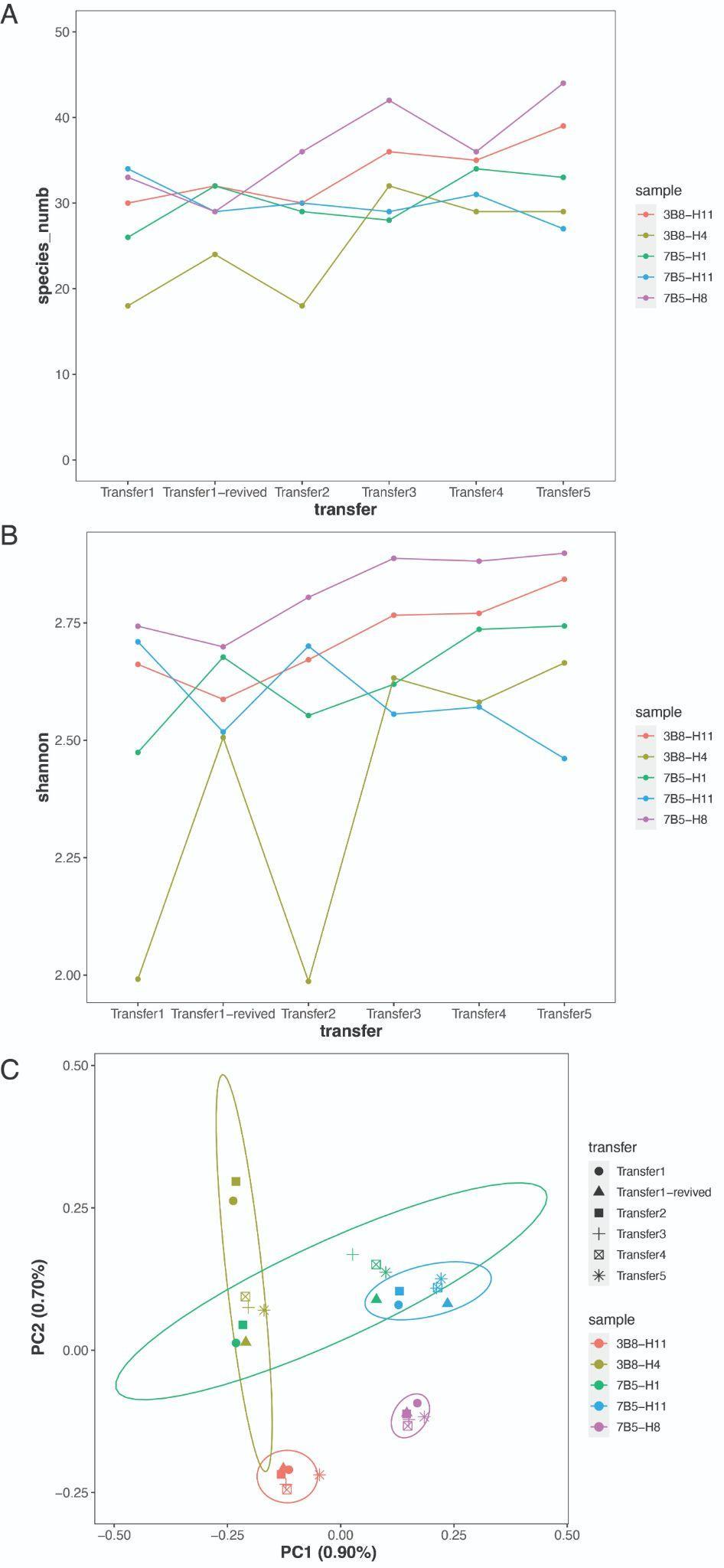

Supplementary Figure S6: Species richness boxplots for samples with different original inoculum and amended with different carbon substrates comparing different generations. Species richness numbers are depicted by dashed lines from all samples using the same original inoculum.

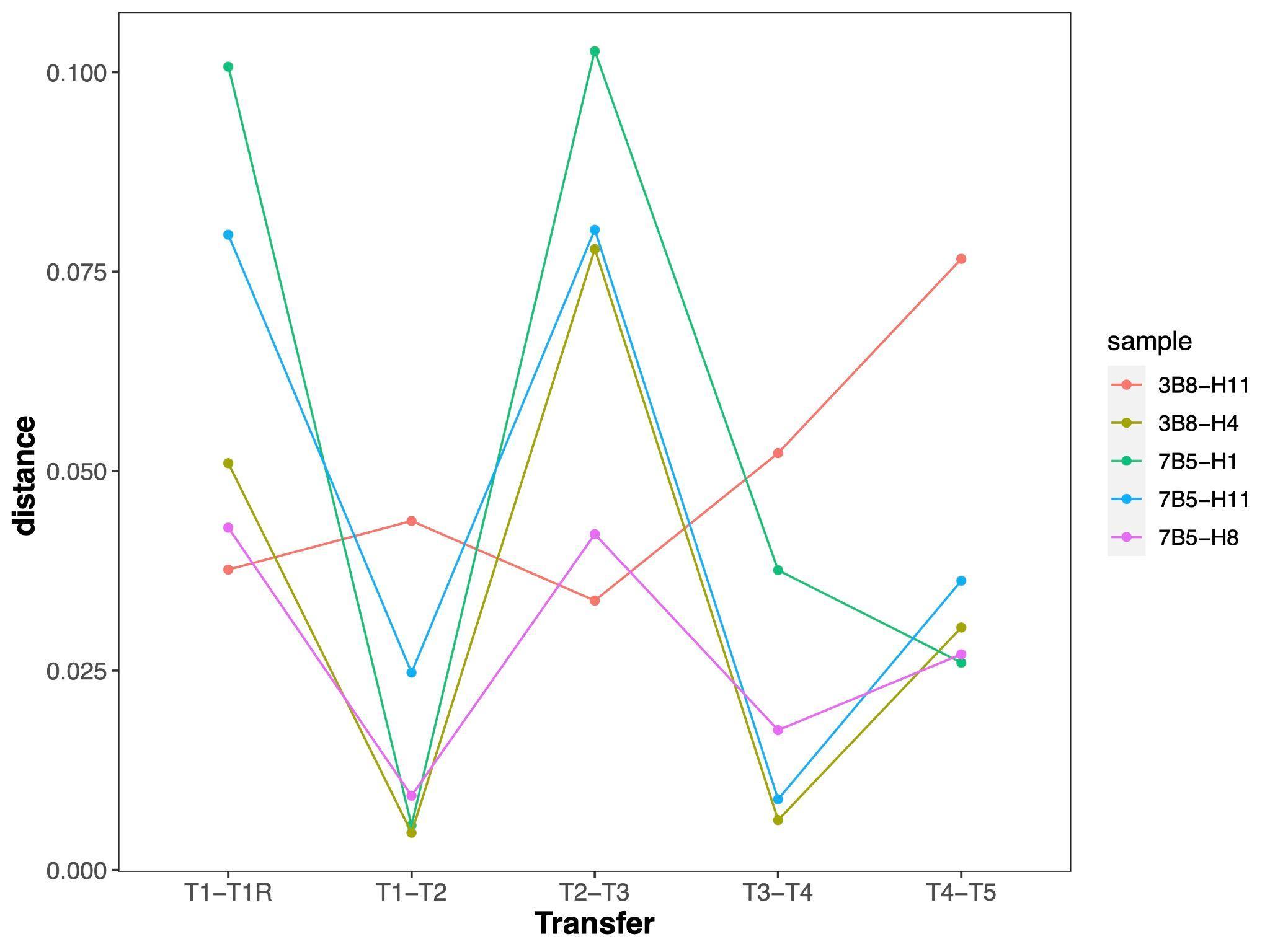

Supplementary Figure S7: Species richness boxplots for samples with different original inoculum and amended with different carbon substrates comparing different generations. Species richness numbers are depicted by dashed lines from all samples using the same original inoculum.

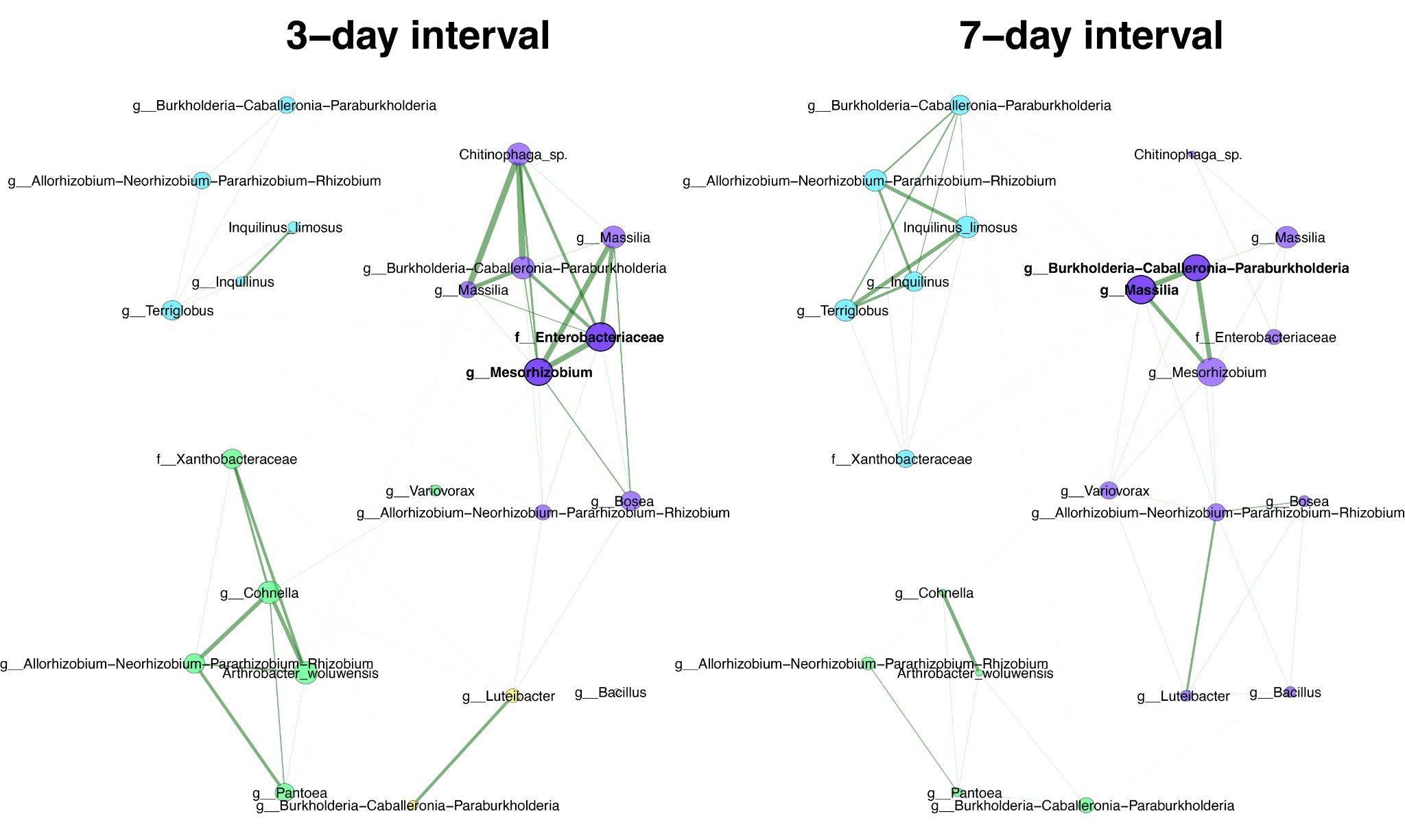

Supplementary Figure S8: Comparison of bacterial associations between derived consortia from 3-day and 7-day transfer intervals. Pearson correlation (>0.3) is used as an association measure, and t-test (< 0.05) is used as a sparse matrix generation method. Eigenvector centrality is used for defining hubs and scaling node sizes. Node colors represent clusters, which are determined using greedy modularity optimization. Clusters have the same color in both networks if they share at least two taxa. Green edges correspond to positive estimated associations and red edges to negative ones. The layout computed for the 3-day interval network is used in both networks. Hubs are highlighted with bold boundaries and corresponding taxon names.
